# Evolutionary rescue of freshwater copepods during historical lake acidification

**DOI:** 10.1101/2025.02.27.640569

**Authors:** Mathilde Salamon, Maxime St-Martin, Rowan Barrett, Alison Derry

## Abstract

The persistence of populations facing severe environmental disturbance can be enabled by natural selection on heritable phenotypic variation - a process known as evolutionary rescue. Few studies have documented this process in complex natural settings and the long-term outcome of evolutionary rescue. Here, we used copepod resting eggs of *Leptodiaptomus minutus* from three time periods of lake ecological history, spanning ≈ 200 generations (100 years) in two populations impacted by historical acidification. Whole genome sequencing of the resting eggs revealed significant allele frequency shifts associated with the acidification followed by pH recovery. We used a resurrection ecology approach to retrace adaptive shifts concomitant with environmental transitions. Copepods from the pre-acidification period showed sensitivity to acidity, while individuals from the acidification period were adapted to acidic pH. This tolerance was subsequently lost during pH recovery, implying an adaptive reversal. Demographic models indicated a decline during the acidification process followed by population recovery based on historical data, suggesting that selection led to evolutionary rescue. This study fills a critical knowledge gap about the long-term implications of evolutionary rescue in the wild.

## Introduction

Predicting adaptive responses to rapid anthropogenic impacts on the environment is an important goal in evolutionary biology [1–3]. The process of **evolutionary rescue** (i.e., an increase in the frequency of adaptive alleles following an initial population decline that allows demographic recovery) has been the focus of many studies because of its potential to allow populations to avoid extinction from sudden environmental degradation [4–10]. Examples of evolutionary rescue have been hinted at in wild populations [8,11]. For instance, field crickets are hypothesized to have escaped high mortality rates from invasive parasitoid flies via a single gene mutation that suppresses the sexual song of males, which attracted the flies [12,13]. However, most studies on evolutionary rescue have been laboratory experiments and theoretical modeling, and clear documentation of this process under natural conditions has been challenging [5]. This may be due to the difficulty of documenting the interplay between demographic, genetic, and environmental factors during an evolutionary rescue event, particularly in complex, natural settings where many ecological and evolutionary processes act concurrently [8]. Although evolutionary rescue has been proposed as a conservation measure for populations facing extinction [14], further evidence of the possibility of evolutionary rescue in nature is needed, particularly because some recent studies have also reported that adaptation can be constrained or even fail to rescue populations [15–17].

An additional knowledge gap is that the long-term consequences of evolutionary rescue in the wild are largely unknown, likely due to the intensive requirements of long-term monitoring of natural populations following an environmental disturbance [8]. Laboratory experiments and observations in the field suggest that it could potentially have detrimental effects over the long term [18]. Indeed, the long-term benefits of evolutionary rescue could potentially be mitigated by two effects: 1) a loss of genetic diversity due to strong selection on adaptive alleles and demographic stochasticity, thereby reducing the ability of populations to adapt to future disturbances [18] and 2) when ecosystems eventually recover or move toward another equilibrium, the phenotypic trait value that was selected for during the disturbance may be distant from the adaptive peak in the restored environmental conditions [8,19–21]. However, environmental recovery may also lead to positive outcomes such as an adaptive reversal [22,23] due to the relaxation of selection [24]. The presence of an adaptive reversal following an evolutionary rescue event would imply that the latter does not necessarily prevent future adaptations[23,25,26]. It is thus critical to study population demography, genetics, and environmental characteristics jointly during and after evolutionary rescue in the wild, as this combination of factors may be especially important in determining the outcome of evolutionary rescue [5].

Resurrection ecology constitutes a powerful toolkit to explore how environmental conditions impact demography and genetics during evolutionary rescue over long time periods [27,28]. Many aquatic organisms produce resting stages as part of their life cycle, which can accumulate in sediments over time and form ‘seed banks’[29]. These resting eggs can be hatched in the lab after being exposed to the proper hatching cues and reared in different environmental conditions in a common garden or reciprocal transplant experimental setup [27,30]. Genomics tools can document (adaptive) allele frequency changes over time [31] and reconstruct demographic history through modeling [32]. With a combination of resurrection ecology and genomics, resting eggs can be used to retrace the evolutionary trajectories of populations relative to environmental disturbances [28,33,34]. The synthesis of these two methods offers a powerful approach to addressing evolutionary questions because it has the potential to illuminate past phenotypic and genetic changes that occurred in a population [27,34].

The ecological history of disturbance and recovery in the lakes of Killarney Park, Ontario (Canada) provides a valuable study system for testing for evolutionary rescue and its potential effects on population resilience in aquatic organisms. Many lakes in this region were acidified in the 1960s as a result of metal mining emissions but have undergone chemical recovery in water pH over the last several decades following government controls on emissions in the 1970s [35]. Lake acidification resulted in high mortality rates and local extinctions of fish [36], zooplankton [37], and phytoplankton [38]. However, a species of a calanoid copepod, *Leptodiaptomus minutus*, developed resistance to acidic pH and subsequently became dominant in zooplankton communities across this acidified landscape [21,37,39]. In contrast, there are examples in which the development of tolerance to acidity did not occur in this species, such as a whole lake acidification experiment in which a population of *L. minutus* crashed until extinction [40]. The high abundance of *L. minutus* during that period [37] hints that populations from the region could have experienced the U-shaped demographic trajectory with recovery due to adaptation that is characteristic of evolutionary rescue.

In the present study, we evaluate the evidence for evolutionary rescue in copepod populations from two pH-recovered lakes with a history of acute acidification. We define evolutionary rescue as a significant initial population decline followed by a population rebound due to the increase in adaptive phenotypes/genotypes frequency [8,9]. Previous research on Killarney Park lakes using resurrected nauplii from resting eggs of *L. minutus* dated from the pre-industrial period (i.e., unexposed to acidity) showed variable survival to acidity in contrast with the consistently high survival of resurrected individuals from the acidification period. However, this was then followed by a return to lower but variable survival to acidic pH during the recovery period [21,41]. If the rise and fall of acid tolerance (indicated by high survival rates and its correlated trait fecundity) represents evolutionary rescue followed by adaptive reversal, we expect that resurrected individuals from the acidification period should have lower fitness at neutral pH than at acidic pH (i.e., a fitness trade-off) and that copepods from the recovery period should have lower fitness at acidic pH. At the genomic level, we expect to observe signatures of directional selection between the pre-acidification and the acute phase of lake acidification. The potential demographic effects of the initial low acid tolerance, combined with the development of acid tolerance over time and the high density of *L. minutus* during the acidification period, hints at the potential U-shaped demographic signature of evolutionary rescue [9]. We should also observe a pattern of reduction in effective population size during the acidification phase followed by a potential rebound indicative of the U-shaped demographic signature of evolutionary rescue. We tested for evidence of the selection of acid-tolerant phenotypes and alleles during the lake acidification, followed by the selection of acid-sensitive phenotypes and alleles during the lake pH recovery, by using resting eggs sampled from sediment cores at three time points of lake ecological history (pre-acidification, acidification, and recovery). Our approach is two-fold: First, we conducted a laboratory reciprocal transplant that employed resurrected *L. minutus* individuals to measure survival, fecundity, and development time at two pH treatments (acidic and neutral pH). Second, we analyzed temporal changes in allele frequencies based on whole genome sequencing (WGS) across the time series. By combining resurrection ecology with WGS, our study expands beyond previous work on the system [21,41] to document the process of evolutionary rescue and its long-term consequences.

## Methods

### 1. Study sites, sampling, and physicochemical data collection

Killarney Park, Ontario, Canada (46°01′ N, 81°24′ W) contains over 600 boreal lakes from the Canadian Shield geological region with pH ranging from 4.3 to 7. This wide range in pH among lakes is the result of natural pH variation, as well as anthropogenically induced acidification associated with the long-range transport of acid deposition from SO_2_ emissions from nearby metal mining smelters during the mid-1900s [35]. Subsequent emission reductions in the mid-1970s led to differences in pH recovery trajectories between lakes due to differences in acid buffering capacities originating from the geological properties of their watersheds [42].

We collected sediment cores from two lakes that acidified and are now at different stages of pH recovery: George (N46°01.7960 W81°23.8310) for which the pH recovered to circumneutral in 1994, and Lumsden (N46°01.5240 W81°25.9730), which is still recovering (Fig. S1B). These two lakes were also impacted differently by the acidification [43], with a lower and more progressive drop in pH for George and a faster, stronger acidification in Lumsden [42]. We sampled 53 sediment cores from each lake during July 2018 and June to July 2019 from the deepest point of each lake. We divided sediment cores into 1 cm-thick sections on-site using a custom-made sediment core extruder and then stored each sediment section in airtight Whirl-pack® bags that were refrigerated in the dark until analysis. We isolated resting eggs of *Leptodiaptomus minutus* from the sediment cores of each lake, representing different periods of ecological history (pre-acidification, acidification, and recovery), with the exception of George Lake, for which we were unable to obtain viable *L. minutus* resting eggs from the pre-acidification period due to low resting egg density. The time periods were defined based on historical pH records for these lakes (see Fig. S1) and pH trajectory reconstruction using diatom and chrysophyte paleofossils [43]

We conducted ^210^Pb dating on one sediment core per lake at the Geotop Center (Université du Québec à Montréal). We then used the rplum package v0.5.1 [44] with the R Statistical Software v4.4.2; [45] to calculate ^210^Pb dates for core intervals (Figure S1A). We used a YSI Pro Plus multiparameter probe in situ (model 10102030; Yellow Springs Inc.) in 2018 and collected water samples in 2019 to obtain physicochemical properties for George and Lumsden lakes (surface temperature, pH, conductivity, dissolved organic carbon, total phosphorus, total nitrogen). Details of the analytical methods for water chemistry and physicochemical parameters of the lakes can be found in the Supplementary Methods and Table S1.

### 2. Resurrection ecology experiment

From the resting eggs recovered from the sediment cores, we applied a resurrection ecology experimental approach in which F_1_-generation copepods were hatched from resting eggs originating from the three time periods and exposed to neutral (pH 6.5) and acidic pH (pH 4.5) to measure survival, fecundity, and development time throughout their lifetime (∼1 month, Fig.1C). These pH values correspond to the circumneutral pH 6-7 at during the pre-industrial and recovery periods in George, and to the recorded acidic pH in Lumsden respectively (Fig. S1). No maternal effects were standardized in this experiment as we were unable to obtain enough hatchlings from the reproduction of the F_1_ individuals to carry on the experiment to the F_2_ generation. Details of the hatching methodology and replication design are provided in the Supplementary Methods and Table S2.

**Figure 1:**
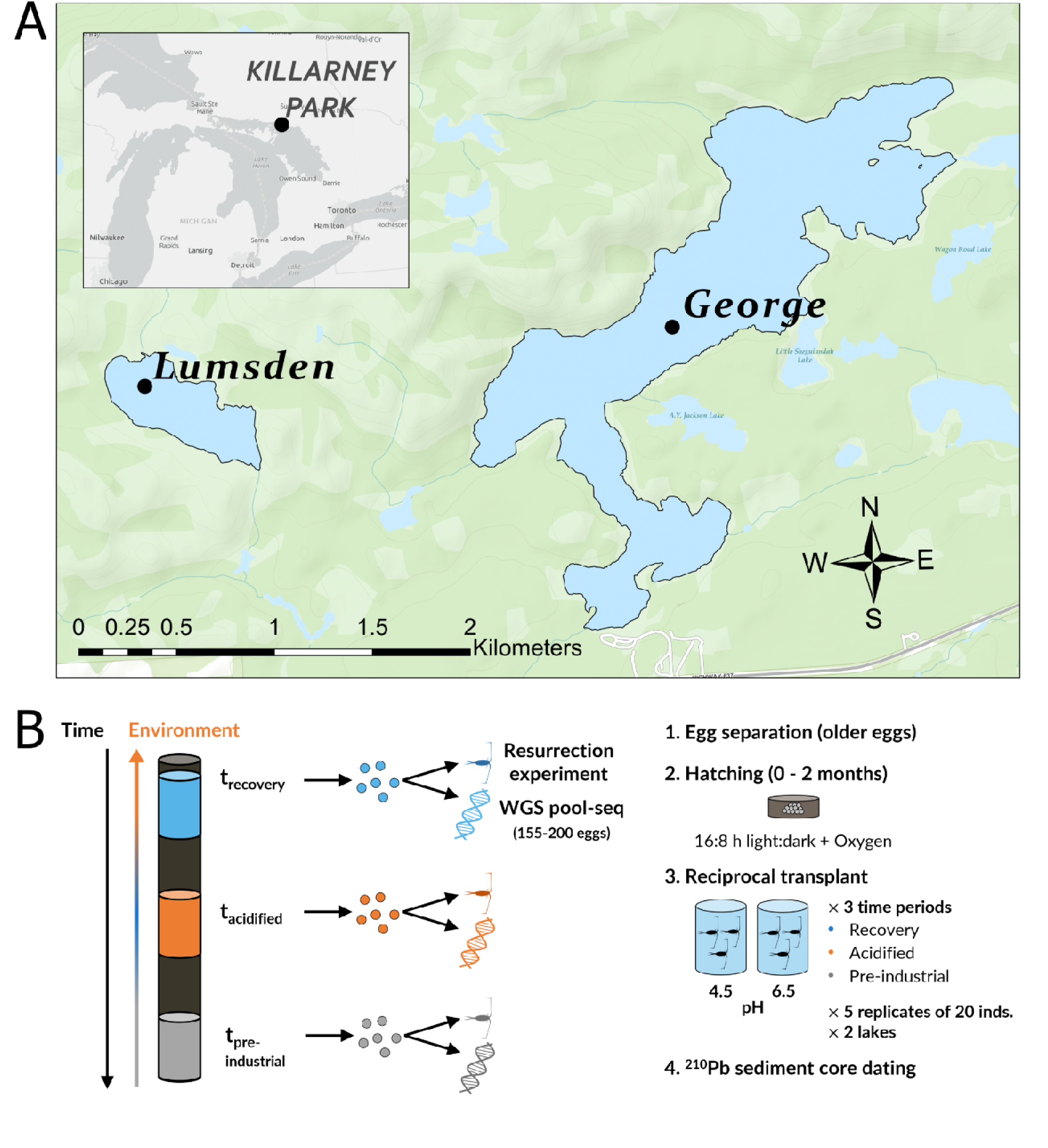
Location of the sampled lakes and experimental design of the resurrection ecology experiment. **(A)** George and Lumsden lakes are located in Killarney Provincial Park ON, Canada. **(B)** Left: Schematic representation of the resurrection experiment and the temporal genomics, right: experimental design.

We observed copepods in each replicate during water changes to record 1) their survival, 2) the number of eggs or directly hatched nauplii produced throughout their lifetime, and 3) the developmental stage (see Supplementary Methods for growth conditions). We measured the survival of copepods by counting the number of living and dead individuals. We obtained the developmental stage of copepods by counting the number of individuals observed at each life stage: nauplii, copepodites, and adult copepods. Their fecundity was measured by counting the number of eggs or nauplii produced relative to the number of adult females present in the culture.

### 3. Analysis of life-history datasets

We analyzed survival, fecundity, and development times with generalized linear mixed effect models (GLMM), generalized linear models (GLM), and t-tests (see supplementary methods). To evaluate the fixed effect on survival in our GLMMs, we used the Akaike information criterion AIC approach corrected for small sample size (AICc) with the ΔAIC criterion from the R package ‘bbmle’ [46]. We kept the effects when ΔAIC > 2. To evaluate the fixed effect on fecundity in our GLMs, we used an analysis of deviance with an F approximation due to the overdispersion. For the development time from Lumsden, we used the likelihood ratio test to examine the fixed effects. The fixed effect coefficients of the GLMs and GLMMs and their confidence intervals were converted to odds ratios (survival), incident rate ratios (fecundity), and fold change (development time) using an exponential function.

### 4. Whole genome sequencing and SNP calling

We applied a pool-seq approach to the sequencing of the resting eggs temporal samples from the two lakes (Lumsden: pre-acidification, acidification, recovery; George: acidification, recovery). The libraries were sequenced on an Illumina NovaSeq6000 Sprime v1.5 to generate 150 bp paired-end reads. As a draft genome for *L. minutus* or a closely related species was unavailable, we applied the reference-free SNP (single nucleotide polymorphism) discovery method from discosnp++ to call SNPs [47]. This generated a dataset of 4,211,276 biallelic SNPs after filtering (see Supplementary Methods for details on the library preparation, Quality Control, SNP calling and filtering).

### 5. Detecting signals of selection

To assess the presence of putative loci under selection during the acidification and pH recovery time periods, we used two approaches: 1) an F_ST_ scan for loci showing exceptional differentiation between time periods, i.e. included within the 95% percentile of the distribution of pairwise F_ST_ values across all temporal comparisons and lakes (top 5%, F_ST_ > 0.29), and 2) an environmental association analysis using the Aux model implemented in Baypass [48]. We conducted the F_ST_ scan approach with the poolfstat R package [49] to detect outlier SNPs showing large differences in allele frequency between time points. The Aux model detected SNPs significantly associated with the acidification (coded as a binary covariable; -1: pre-acidification and recovery, 1: acidification) in both the Lumsden and George datasets. We considered SNPs to be significantly associated with the acidification covariable when their Bayes Factor obtained from the Aux model (BF_mc_) exceeded > 20 deciban units (dB) [48].

### 6. Population structure and demography

We first estimated population structure with the core model from Baypass, then compared the results to a genome-wide pairwise F_ST_ matrix estimated with poolfstat. We also estimated the observed heterozygosity for each temporal sample with poolfstat. To assess the possibility of a demographic decline due to the acidification event (i.e. the first phase of the hallmark U-shaped trajectory of evolutionary rescue) we used the diffusion approximation approach from δaδi to identify a potential decrease in effective population size N_e_ [50]. We selected the recovery temporal samples of Lumsden and George for our dataset and analyzed each population separately (1D models). The most complex model (“bottlegrowth”) has a U-shaped demography as described above, with a bottleneck starting at the time T_B_ (known scaled time in units of 2N*_ref_* X generations between the acidification peak and present), followed by effective population size recovery (Fig. 3A), we also analyzed three simpler models, including two nested models: a) a genetic bottleneck without recovery (“two epochs” model), b) N_e_ growth, and c) neutral (Fig. 3C and S3). More details are provided in the supplementary methods.

## Results

### 1. Resurrection ecology experiment: Adaptive shifts in life-history traits following acidification and pH recovery

The resurrection ecology experiment revealed population life history traits shifts (lifetime survival, development time, and fecundity) that were consistent with the adaptation of *L. minutus* to lake acidification, followed by a reversal during recovery (Fig. 2, Tables S2 and S3). For the survival of copepods from nauplii to adulthood (lifetime survival) in both Lumsden and George lakes (Fig. 2A), we found significantly greater survival rates of the resurrected individuals from the acidification period than from the recovery period. Specifically, for Lumsden, there were no significant differences in survival between individuals from the pre-acidification and recovery periods. These results held regardless of pH (non-significant interaction). Although this interaction was not significant, survival was also significantly higher at neutral pH compared to acidic pH. We found similar results, albeit less pronounced, for early survival (from hatching into nauplii to metamorphosis into copepodid), except for the effect of pH, which was not significant (Fig. S4A, Tables S2 and S3). Copepods from the acidification period developed significantly faster to adulthood compared to the pre-acidification period for Lumsden, and for both Lumsden and George, there was no difference in the development time between acidification and recovery individuals (Fig. 2C, Tables S2 and S3). The pH treatment and the interaction between time period and pH did not affect development time to adulthood for both lakes (Table 1). The results were similar for early development (Fig. S4B) except for a significant difference between acidification and recovery for Lumsden (Tables S2 and S3). Female copepods from Lumsden were significantly more fecund during the acidification period than during the pre-acidification and recovery periods, but the pH and the interaction between time period and pH did not affect their fecundity (Fig. 2B, Tables S2 and S3). However, George Lake female fecundity was only marginally affected by time period and the time period X pH interaction. To summarize, our experiment suggests there was an acquisition of acid tolerance during the acidification period followed by a loss of tolerance, shown through differences in survival and fecundity. The increase in acid tolerance was accompanied by faster development, but this change in development rate did not reverse to pre-acidification levels following pH recovery. These results were consistent across both lakes despite having experienced different strengths of historical acidification (Fig. S1).

**Figure 2:**
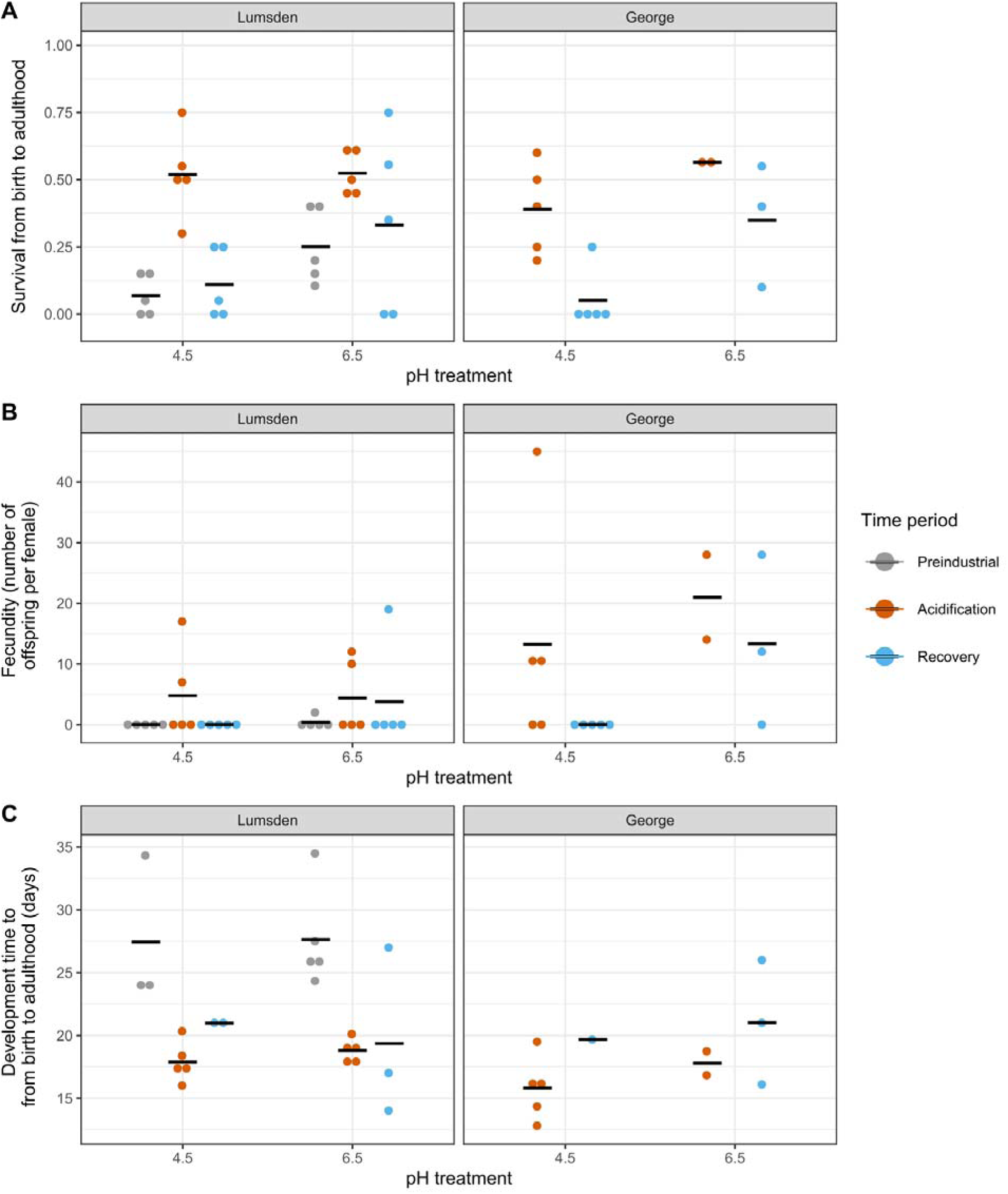
Observed shifts in life-history traits in freshwater copepods from different ecological periods (pre-acidification, acidification, recovery) in two historically acidified lakes as a response to two levels of pH treatment as revealed by the resurrection ecology experiment. **(A)** Lifetime survival, **(B)** fecundity (number of eggs or nauplii per female) and **(C)** development time from birth to adulthood. Each dot represents a measurement for one replicate, and the black bars indicate the mean response for each treatment.

### 2. Population genomics: demography, population structure, and rapid genetic adaptation to acidification and lake pH recovery

Using the recovery time period samples from George and Lumsden lakes, we found indications of significant declines in their effective population sizes during the acidification period. The best models based on the highest likelihood for both populations were the models including population contraction at a known fixed time in the past corresponding to the acidification event. (Fig. 3, Table S5). Figure 3 (panel C and D) show the folded Site Frequency Spectrum (SFS) for the observed data and the model, which correspond to a histogram of the number of SNPs as a function of the allele count in the population. The SFS is folded (i.e. only focuses on the minor allele), as ancestral alleles are unknown. For Lumsden Lake, the likelihood ratio test led to the rejection of the nested model of growth (Fig. S5, Table S5), favoring the more complex model of bottleneck followed by growth (Figure 3A and B, Table S5, adjusted D-statistic = 5.29, p-value = 0.02), whereas the nested growth model was favored for George Lake (Figure 3C and D, adjusted D-statistic = 1.12, p-value = 0.29). For the latter, the small *Nu* parameter (ratio of contemporary effective population size to ancestral effective population size *N_ref_*) value obtained for the growth model implies a population decline toward a very small effective population size (Table S5). Our analyses also suggest that the ancestral effective population sizes were large, with estimated θ values of 2.9 X 10^6^ for George and 1.5 X 10^6^ for Lumsden (see Table S5; 0 = 4N_ref_µL, with *N_ref_* the ancestral effective population sizes, μ the mutation rate, and *L* the effective sequenced length). In contrast, the decrease in effective population size was likely very pronounced, with very low inferred values for the ratio *Nu* in George Lake (3.5 X 10^-5^, Table S5), and for the ratio *NuB* of the effective population size after the genetic bottleneck to the ancient effective population size *N_ref_* in Lumsden Lake (2.3 X 10^-5^, Table S5). These parameter values should be interpreted with caution given the high uncertainty around the parameter estimates (Table S5). Demographic decline was accompanied by a decrease in heterozygosity in Lumsden from 0.30 to 0.28, followed by a smaller increase to 0.29 during the recovery phase. Conversely, for George, we did not find a change in heterozygosity during the recovery (observed heterozygosity = 0.28 for both the acidification and recovery time periods). We found that pairwise F_ST_ values were higher between lakes across time periods (moderate, 0.09 on average) than between time periods within lakes (low, 0.03 on average). For Lumsden, genetic differentiation was slightly lower between the pre-acidification and the recovery periods than between the acidification and the pre-acidification or recovery (Table S6).

**Figure 3:**
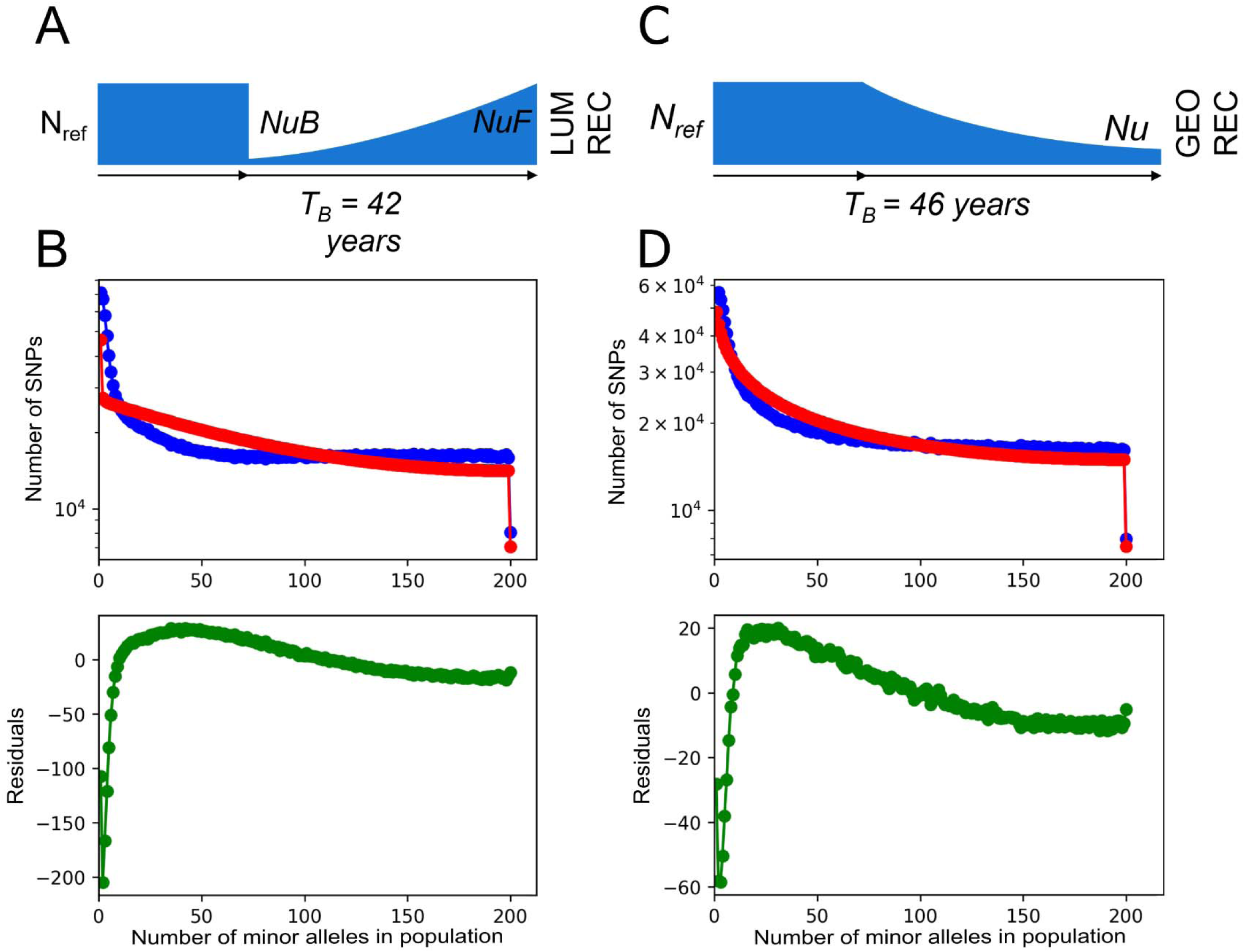
Results of the best demographic model tested with δaδi. **(A)** Visual 1D demographic model of Lumsden Lake, with a genetic bottleneck at the time of acidification, followed by exponential recovery. *NuB* and *NuF* are the ratio of effective population sizes after the bottleneck and the recovery to the ancient effective population size *N_ref_*., respectively. T_B_ is the known scaled time (in units of 2N*_ref_* X generations) between the acidification peak and present **(B)** Top: Folded site frequency spectrum (SFS) for a sample size of 400 (two times the number of individuals) for the observed data (blue line) and the model (red line), showing the logarithm of the number of SNPs (y-axis) as a function of a given read count (x-axis). Bottom: residuals of the normalized differences between the observed data and the model. **(C)** Visual 1D demographic model of George Lake, with a growth event starting at the time of acidification T_B_. *Nu* is the ratio of contemporary effective population size to ancestral *N_ref_*. **(D)** Folded SFS (top) and residuals between the observed data and the model for George Lake (bottom).

The genome scan of pairwise F_ST_ values detected outlier loci in temporal comparisons of both study lakes (Lumsden: pre-acidification vs acidification and acidification vs recovery, George: acidification vs recovery). The 95% percentile (top 5%) of pairwise F_ST_ values across all time comparisons for both lakes was F_ST_ > 0.29, which was used as a threshold to detect outlier SNPs. For Lumsden, we found 165,087 outlier SNPs in the pre-acidification vs acidification comparison (≈ 4% of the dataset), and 232,515 outliers in the acidification vs recovery comparison (≈ 6% of the dataset). For George, we found 157,597 outlier SNPs in the acidification vs recovery comparison (≈ 4% of the dataset). We also found 19,525 outlier SNPs in common between the two-time comparisons for Lumsden in the F_ST_ genome scan and significantly associated with the acidification in the Baypass Aux model (BF_mc_ > 20). Remarkably, all of these outliers showed a reversal in allele frequencies during the recovery period (Fig. 4A). These SNPs are considered candidates for directional selection during the acidification followed by an adaptive reversal during recovery. We identified 78,864 SNPs that were outliers in the Lumsden F_ST_ scan for the pre-acidification to acidification comparison, but not for the acidification to recovery comparison, and not significantly associated with the acidification in Baypass (BF_mc_ < 20, ≈ 2% of the dataset, Fig. 4B). The remaining 141,365 Lumsden outlier SNPs were not significantly differentiated in the F_ST_ scan for the pre-acidification to the acidification comparison, but were significant for the acidification to recovery comparison, and not significant in Baypass (BF_mc_ < 20, ≈ 3% of the dataset, Fig. 4C). We also found 5,907 outlier SNPs in common in the F_ST_ scan for the acidification to recovery temporal comparisons in both Lumsden and George lakes and significant in Baypass (BF_mc_ > 20, ≈ 0.1% of the dataset, 1% of the total outliers across methods), and 100% of these outliers had allele frequency changes in the same direction for this time period transition. These SNPs are considered candidates for parallel selection from the increase in pH during the recovery phase.

**Figure 4:**
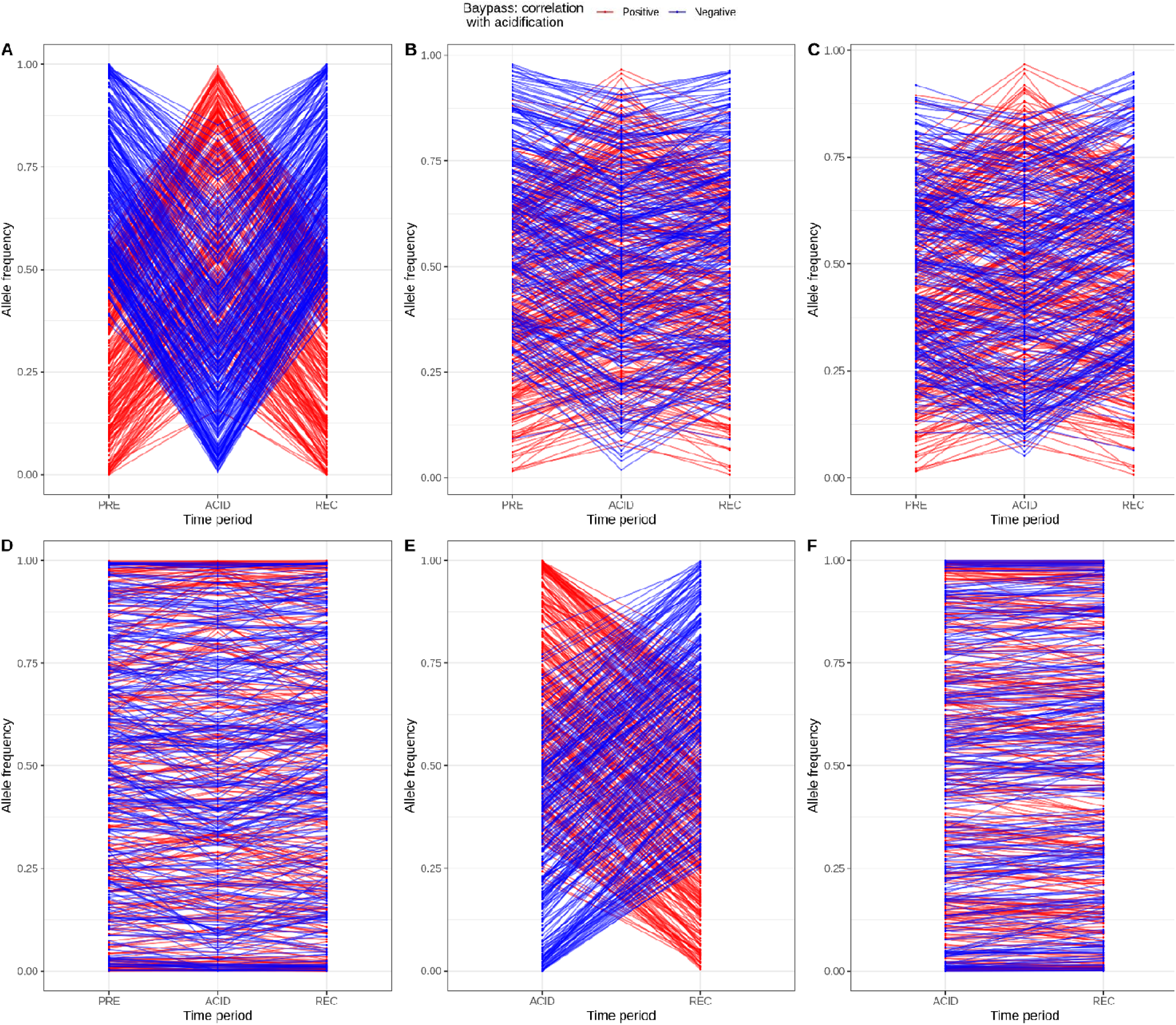
Allele frequency changes of outliers detected with the F_ST_ scan and Baypass. SNPs are significantly associated with lake acidification in the Baypass Aux model if Bayes Factor BF_mc_ > 20 dB, and not associated when BF_mc_ < 20 dB. The red and blue colors indicate positive and negative associations with acidification in Baypass (regression coefficient of the association between the SNP allele frequencies and the covariable β_ik_ > 0 and β_ik_ < 0 respectively). Plots show a random subset of 500 SNPs from the category of outlier and non-outlier SNPs described for Lumsden and George Lake copepods. **(A)** Lumsden SNPs significant in the F_ST_ scan for both temporal comparisons and significant in Baypass, **(B)** Lumsden SNPs significant in the F_ST_ scan for the pre-acidification to acidification comparison but not for the acidification to recovery comparison, and not significant in Baypass, **(C)** Lumsden SNPs not significant in the F_ST_ scan for the pre-acidification to acidification comparison but significant for the acidification to recovery comparison, and not significant in Baypass, **(D)** Lumsden non-outlier SNPs: non-significant in the F_ST_ scan for both time comparisons and not significant in Baypass, **(E)** George outlier SNPs: significant in the F_ST_ scan and Baypass **(F)** George non-outlier SNPs: non-significant in the F_ST_ scan and Baypass.

Using the core model in Baypass as our outlier analysis, we found 36,110 outlier SNPs with p-value < 0.001, which represented ≈1% of the dataset (Fig. S6 and S7). The XtX estimate (equivalent to SNP-specific F_ST_) is based on the variance in allele frequencies across populations, thus, in the present case, outlier SNPs are significantly differentiated considering both the populations of origin (Lumsden and George lakes) and temporal samples. For the Aux model testing the association between SNP allele frequencies and acidification, we found 176,744 outlier SNPs significantly associated with acidification (BF_mc_ > 20; ≈ 4% of the dataset). These outlier SNPs have allele frequencies both negatively and positively associated with acidification (Fig. 4, Fig. S8), and some showed a strong association, with their correlation coefficient of the association between the SNP allele frequencies and the covariable |β_ik_| > 0.2 (33,279 outliers, Fig. S8). Finally, we found outliers in common between the F_ST_ scan and the Aux model in Baypass (Fig. S7): 64,767 outliers for Lumsden (using the SNPs that are outliers in the F_ST_ scan in at least one temporal comparison) and 43,249 outliers for George (Fig. 4E).

## Discussion

Providing evidence of evolutionary rescue and documenting its long-term consequences in complex natural ecosystems is crucial for predicting populations’ responses to ongoing environmental change [8]. Here, we investigated the possibility of evolutionary rescue in freshwater calanoid copepod (*Leptodiaptomus minutus*) populations that were historically exposed to acute lake acidification. We combined a resurrection ecology experiment with whole-genome data spanning ≈ 100 years (200 generations) in two lakes with histories of acidification. We found adaptive shifts during acidification, with increased acid tolerance and fecundity as well as faster growth rate, which were probably mediated by directional selection at the genomic level. We also found genomic evidence for a population decline from lake acidification followed by historically documented high population density, suggestive of evolutionary rescue. Resurrected copepods from the recovery period showed reduced acid tolerance and fecundity, accompanied by a reversal in allele frequency at candidate loci. Overall, our results provide evidence in support of evolutionary rescue from lake acidification in these wild populations, followed by an adaptive reversal during pH recovery.

### 1. Adaptation to acute historical acidification

We detected shifts in life-history traits between the pre-acidification and acidification time periods for copepods from Lumsden Lake. We documented an increase in lifetime survival and fecundity as well as faster development rates overall (across pH treatments) for individuals from the acidification period. Similar trait values were also observed in individuals from George Lake for the same time period, although we were unable to compare with the pre-acidification period for that population. As our resurrection experiment was conducted within a single generation and thus could not account for maternal effects, we acknowledge that the experimental setup did not allow us to disentangle the influence of genetic versus plastic and maternal effects on the life-history traits. We also could not control for the effect of using different hatching methods for the recovery period compared to the acidification and pre-acidification periods, which could correlate with the phenotypic traits measured. For example, it is possible that direct hatching without egg isolation and a period of storage could result in higher fitness. However, copepods from the acidification period had the highest survival and fecundity compared to the other two time periods studied, and they also had different trait values than copepods from the pre-acidification period despite using the same hatching methods. Apart from survival, for which the effect of pH was significant, fecundity and development time showed flat reaction norms across pH treatments, with most differences occurring between individuals from distinct time periods. Considering that the individuals from different time periods correspond to distinct genotypes within the same continuous population (albeit there is a possibility of migration, see below), this implies that we did not find indications of genotype X environment interactions for these traits. This suggests that plasticity likely plays a limited role in changes in fecundity and development as a response to acidity. Fecundity was low overall at pH 6.5, which could be due to the temperature and light regime used for the experiment, or the use of an artificial medium (COMBO) instead of lake water. It is unlikely to have resulted from using inappropriate food, as both selected algal species are good food sources for calanoid copepods [51]. There were some signs of initial intraspecific variation in acid tolerance, as a few individuals from the pre-acidification period from Lumsden Lake were able to survive at acidic pH until sexual maturity. These acid-tolerant phenotypes could have been positively selected during the process of acidification, leading to adaptation to acidic pH. Our experiment also revealed an important concomitant decrease in development time following the acidification for both early and late development stages. These results are consistent with adaptation to historical lake acidification leading to a faster pace of life, with higher survival, fecundity, and increased growth rates, which have also been reported in response to stressful environments, such as urbanization [52].

The adaptive shifts in life-history traits that we detected were accompanied by significant allele frequency changes between the pre-acidification and the acidification periods. This implies that the life-history shifts observed between these two time periods could be due in part to shifts in adaptive allele frequency, resulting from directional selection due to acidification. As these allele frequency changes were both positively and negatively associated with lake acidification, and we do not know the ancestral allele states, adaptation could have occurred through positive selection, purifying selection, or a combination of both [53,54]. The high number of outlier SNPs detected with our two genome scan analyses also suggests that linked selection could have played a role in structuring genomic responses in these populations [55]. *Leptodiaptomus minutus* does not currently have an annotated genome, and thus we were unable to conduct functional analyses of the SNPs putatively under selection from acidification. Possible functions under selection to explore in future studies could be related to osmoregulation and homeostasis, as these physiological functions have consistently been identified as a mechanism of response to acidification in copepods [56,57]. Regulation of diapause [58,59] and lipid metabolism [60] are also of interest as they tend to be favored in stressful environments. Taken together, the changes in life-history traits (survival, fecundity, and development), as well as significant shifts in the frequency of putatively adaptive alleles, provide support for the occurrence of rapid evolution due to historical changes in pH.

### 2. Demographic consequences of acidification and evolutionary rescue

The universal increase in survival, fecundity, and growth rates that we observed in copepods that experienced lake acidification suggests that individuals from this time period have higher fitness than individuals from the pre-acidification period. During the initial process of acidification, the lower survival and fecundity at acidic pH, combined with a slower development rate, likely had important demographic effects, potentially triggering a population decline. Our demographic analyses support this possibility, with the best models indicating a genetic bottleneck or a decline in effective population size (N_e_) at the peak of acidification. Population size was probably large initially, as the model parameters suggest a high ancestral effective population size prior to the acidification. The populations from both of our study lakes showed generally similar responses despite different trajectories during acidification (faster and steeper for Lumsden Lake) [43]. These results confirm that the rate of acidification represented a stressful environmental event for aquatic organisms, resulting in a genetic bottleneck or decline in effective population size of *L. minutus* despite it being one of the most tolerant zooplankton species in this region [37,41]. This decline resulted in a moderate reduction in genetic diversity during the acidification period for Lumsden. The population decline due to acidification was followed by a population rebound, as *L. minutus* populations were documented to be at high density in acidified lakes by 1972-1973 [37]. The genetic bottleneck and decline in effective population size detected in our study populations during the acidification, in conjunction with a probable population recovery and an increase in the frequency of adaptive phenotypes and putative alleles, indicate that these populations likely avoided local extinction [61] through evolutionary rescue. Our study provides a rare example in which a probable case of evolutionary rescue has been documented in nature. However, the limited replication (two lakes, one lacking eggs from the pre-acidified period), and the absence of non-acidified control lakes in our study design, prevents us from drawing broad conclusions regarding the repeatability of this process in other populations, or the factors that influenced the rescue.

It is possible that migration played a positive role in facilitating evolutionary rescue [62], since George and Lumsden lakes are embedded in a landscape of nearby lakes. However, it is unlikely that an extinction of the original populations was followed by seeding from surrounding populations. In the case of a local extinction followed by re-colonization, we would have expected to see a pairwise F_ST_ value change between pre-acidification to acidification for Lumsden on the same order of magnitude as the pairwise F_ST_ between Lumsden and George lakes for the same time periods. Indeed, the two lakes are only separated by ∼ 1 km, a similar distance to other surrounding lakes. However, the observed pairwise F_ST_ was at least 2-3 times higher between lakes than within lakes between time periods (Table S6 and Figure S2). Knowledge of gene flow in lake copepods is still limited, and species dependent [63,64] and the potential role of migration on the evolutionary rescue of populations from the Killarney study system will require further investigation.

Despite evidence for evolutionary rescue in *L. minutus* populations, our demographic analyses highlight the potential negative consequences of this process. Indeed, the strong selection that occurred during evolutionary rescue appears to be associated with a genetic bottleneck in the Lumsden Lake population and a decline in N_e_ in the population of George Lake. Even though N_e_ seems to have recovered in Lumsden Lake during the following environmental recovery and increased population abundance, the best model for George suggests that N_e_ did not recover in this population. This is in line with a previous study that found that evolutionary rescue led to a reduction of genetic diversity due to strong selection and drift occurring at reduced population sizes [18]. Prior experimental studies of evolutionary rescue have also shown demographic and genetic bottlenecks with large reductions in N_e_, followed by limited N_e_ recovery [6]. This suggests that the short-term benefits of evolutionary rescue might be accompanied by detrimental effects on genetic diversity over the long term.

### 3. Evidence for an adaptive reversal during lake pH recovery

The results of our resurrection experiment indicated a loss of acid tolerance (survival and fecundity) in individuals from the recovery period for both lakes. In the case of an adaptive reversal, we might expect to find a trade-off between survival under acid stress and survival or fecundity at neutral pH for the copepods from the acidification period, for example, due to antagonistic pleiotropy [24,65]. This would lead to the selection of acid-sensitive phenotypes during pH recovery in the lakes. This pattern has been previously shown in populations of *L. minutus* from acidic lakes and ponds [21,66]. Surprisingly, we did not find indications of such a trade-off in individuals from the acidification period, which showed high fitness across neutral and acidic pH treatments. This pattern could be due to unmeasured maternal effects that had positive effects on copepod fitness from the acidification period, possibly masking the expected trade-off. Additionally, our results are based on a laboratory experiment in a simplified environment, whereas *L. minutus* fitness might have been affected by ecological interactions such as increased competition and predation during the progressive ecological and chemical recovery [39]. One aspect of our experimental design that limits the ability to assess a potential fitness trade-off is that copepods were fed ad libitum. The measured resource allocation traits (fecundity and survival) may be in genetic covariance with traits related to resource acquisition (e.g. body size), and these are unlikely to be selected for if resources are very abundant [67]. Finally, fitness trade-offs from local adaptation are not always detected even if present [68]. This might be the case here due to the complexity of the study system and its inherent stochasticity, which could have masked these effects.

Despite the apparent absence of a fitness trade-off for acid tolerance, we did find signs of an evolutionary reversal at the genomic level. A large number of Lumsden outliers which were significant for both temporal comparisons in the F_ST_ scan (pre-acidification to acidification and acidification to recovery) were also significant in the Baypass Aux model, and all of these outliers showed a reversal in the direction of allele frequency change (either positive or negative) during recovery. Our results thus indicate that the reversal in acid tolerance at the phenotypic level is at least in part due to a reversal in adaptive allele frequency at the genomic level. A similar adaptive reversal at both the phenotypic and genomic levels has previously been shown in *Daphnia magna* in response to shifts in predation levels [23] and for resistant phenotypes of *Daphnia galatea* to toxic cyanobacteria as a food source [22]. In contrast to survival and fecundity, development time showed no sign of reverting to pre-acidification trait values or being responsive to pH. Our results thus indicate that the reversal of life-history traits (fecundity, survival) is due in part to directional selection, whereas development time may be responding to other sources of selection and therefore did not experience a reversal.

### 4. Consequences of evolutionary rescue and future trends

We found a lower proportion of parallel SNPs at the genomic level (1% of the outliers) between George and Lumsden lakes compared to what has been found in other study systems [69]. Our results suggest that both populations went through a similar pattern of evolutionary rescue followed by an adaptive reversal, implying that these processes may have been repeated in other *L. minutus* populations with similar ecological histories of lake acidification and pH recovery. Adaptive reversals have been documented in populations from other historically acidified lakes in the region [21]. These processes might not be highly repeatable at the SNP level, but they could be repeated at the gene or functional level due to redundancy [69].

Although *Leptodiaptomus minutus* populations were sufficiently resilient to undergo evolutionary rescue and adaptive reversals from historical acidification and lake pH recovery, this might not be the case under future threats. Acidified lakes in the Killarney, Ontario (Canada) region have not reverted to their original ecological state [39], and additional stressors relevant to copepod biology appeared during the same time period as the acidification. A potential selection pressure that arose before the acidification period and continued during the pH recovery period is climate change [70]. The reduced effective population size found in one of the *L. minutus* populations suggests that their present and future adaptive potential could potentially be limited [71]. Decreases in *L. minutus* populations were recently reported across several lake ecosystems (Adirondack Park and Killarney Provincial Park) along with ongoing physicochemical and biotic changes [72,73], highlighting the need for an assessment of the adaptive potential to climate change and other stressors in ecologically significant zooplankton, such as freshwater calanoid copepods.

## Conclusions

Our study fills a critical knowledge gap concerning the process of evolutionary rescue in nature. We assessed the long-term consequences of evolutionary rescue and show that this process did not prevent an adaptive reversal during environmental recovery, implying that the loss of genetic diversity that occurs during evolutionary rescue does not necessarily prevent future adaptation.

## Author contributions

MS, AMD, and RDHB designed the study. MS, MSM and AMD collected the field samples. MS and MSM conducted the laboratory experiment. MS performed the DNA extractions, generated the pool-seq processing pipeline, and ran the bioinformatic and statistical analyses, with input from AMD and RDHB. The article was written by MS, with input from AMD and RDHB. All authors contributed to the review and editing. AMD and RDHB funded the study.

## Acknowledgments

This research was supported by a team grant awarded by NSERC Discovery grants to AMD (RGPIN-2016-05143) and RDHB (RGPIN-2019-04549), a QCBS seed grant to AMD and RDHB, and a Canada Research Chair to RDHB. The NSERC Create ÉcoLac Training program, a recruitment award, and doctoral support scholarships from the Faculty of Science of Université du Québec à Montréal provided scholarship funds to MS. We acknowledge financial support from the Groupe de recherche interuniversitaire en limnologie (GRIL), a strategic cluster of the FRQNT. Analyses were performed on the clusters of the Digital Research Alliance of Canada supported by a RAC application to RDHB. We thank Daphnée Trépanier-Leroux, Simon Thibodeau, and Louis Astorg for field assistance. We thank Mathieu Gauthier for support with the software Baypass, Ryan Guntenkunst for the δaδi analyses and Antonin Prijac for the 210Pb dating analyses.

## Data Accessibility

The raw sequencing reads have been deposited in the National Center for Biotechnology Information Sequence Read Archive SRA repository (BioProject PRJNA1229077 https://www.ncbi.nlm.nih.gov/bioproject/PRJNA1229077) and the accompanying metadata are also stored in the SRA (BioProject PRJNA1229077), using the Eukaryotic water MIxS package. The scripts and input data have been uploaded to the Dryad Digital Repository (10.5061/dryad.qbzkh18v6). We included the population used as an anchor on the Dryad repository. The code, raw data and processed results are stored on Zenodo and appear in the Dryad repository under “Software files available” and “Supplemental files available.”

