## Supplementary Material for "Evolutionary rescue of freshwater copepods during historical lake acidification"

**Supplementary Methods**

### 1. Study sites, sampling, and physicochemical data collection

For each sediment core section, we weighed the sediments before (wet weight) and after drying (dry weight) to obtain the dry bulk density *ρ*. For each section, 100 mg of dry weight was isolated, and alpha-particle radiation activity was counted for 48 hours in an alpha counter. To calculate the ^210^Pb dates with the rplum R package, we set the decimal date to 2019.477, with 5,000 simulations and retaining three levels of supported ^210^Pb activity. ^210^Pb dates output by rplum for the sediment cores collected from each lake and the temporal sections and selected for the resurrection and genomics experiments are presented in Figure S1A.

The pH, specific conductivity (SPC; µS/cm and temperature (°C) were measured at 0.5 m depth (Table S1) and throughout the water column (unpublished) in August 2018 with a YSI pro plus multiparameter probe (model 10102030; Yellow Springs Inc.). Other physicochemical parameters were analyzed in the GRIL-UQAM water chemistry laboratory on water samples collected at 0.5 m depth in late June 2019. Dissolved organic carbon concentration was measured from water filtrates passed through 0.45 µm filters (surfactant-free membrane filters) and then acidified (5% phosphoric acid), followed by sodium persulfate oxidation and concentration reading with a 1010 total organic carbon analyzer (O.I. Analytical, College Station, TX, U.S.A.). Total phosphorus was measured spectrophotometrically on the same instrument by the molybdenum blue method after persulfate digestion (Griesbach and Peters, 1991). Total nitrogen samples were also analyzed on this instrument with the continuous flow analyzer by the alkali persulfate digestion method and then coupled to a cadmium reactor, following a standard protocol (Patton and Kryskalla, 2003).

1. Resurrection ecology experiment

We used two different methods to hatch the resting eggs, as the older eggs (pre-acidification and acidification periods) could not be hatched directly [1]. The first resting eggs hatching method was implemented for the eggs corresponding to the recovery period (from the past 10-20 years). Following the method in Hairston & Kearns [2], we added the equivalent of 1 cm of the core sediment to the bottom of a plastic tray and then covered the eggs with 1-2 cm of COMBO culture medium [3] prepared in the laboratory and pH adjusted to 4.5 or 6.5 with 0.02M sulfuric acid. In the second method, we used the eggs from to the pre-acidification and acidification periods, which we separated from the sediments for each time period with a sugar flotation method modified from [4]. We then stored the isolated resting eggs in distilled water in a dark refrigerator at 4°C. Prior to exposure to hatching signals (luminosity, temperature, and oxygenation), we stored the eggs from the acidification period for 2 weeks to 1 month under the above conditions and the eggs from the pre-acidification period for 2 months following Derry et al. [5]. This is necessary for resting eggs that have been buried for longer periods in lake history, as more time is required to stimulate them to hatch [2]. After these storage periods, we transferred the resting eggs from the pre-acidification and acidification time period to a hatching beaker containing 200 mL of COMBO medium. For both methods, resting eggs from several sediment core sections had to be combined in the hatching trays and beakers to obtain enough nauplii for the resurrection experiment, but we did not combine sediments from different time periods or lakes (see table S2 for the number of cores used per lake and time period). There was no technical replicate for the sediment pools used for the hatching nor of hatching trays/beakers. We exposed the resting eggs to the following hatching signals: 16h:8h light:dark cycle, 20°C, and oxygenation [5,6]. We kept the hatching trays and beakers in a Thermo Scientific Precision Model 818 Incubator growth chamber and examined them regularly for the presence of nauplii. For both hatching methods, we started a replicate when there was enough nauplii hatched, aiming for N = 20 nauplii and five replicate beaker per lake, time period, and pH treatment. This was not achieved for George Lake due to the lower resting egg density. We verified the identity of nauplii using the list of species present in our study lakes [7] as well as a key from the Department of Biological Sciences at the University of New Hampshire [8]. We removed a few nauplii from the beakers and datasets that were misidentified (N = 5 cyclopoid copepods and N = 1 calanoid copepod: *Epischura lacustris*).

We transferred hatchling nauplii (≈ 20 individuals per replicate) to beakers with 200 ml of COMBO medium adjusted to either an acidic pH of 4.5 or a circumneutral pH of 6.5. We changed the culture medium regularly (once to three times a week). We fed the copepods with a mixture of 1 mL of brown algae (Cryptomonas sp. CPCC 336) and 1 mL of green algae (Chlamydomonas sp. CPCC 243) cultured at a stationary phase. We cultured the green algae in a modified Bold's Basal medium (Stein, 1973), and the brown algae in COMBO adjusted to pH 6.5, in the growth chambers containing the copepod cultures with a 16h:8h light:dark cycle and at 20°C. We noticed algal growth in the copepod cultures after 2 days, indicating sufficient food abundance.

2. Analysis of life-history datasets

We modeled survival for Lumsden and George lakes with GLMMs using a binomial distribution and logit link function. We modeled fecundity with GLMs using a quasi-Poisson distribution to account for overdispersion. For development time, as the number of replicates was low for George Lake, we used different statistical tests for the two lakes. For Lumsden, we used a GLM with a Gamma distribution and a log link function, and for George, a t-test pooling data over the pH treatment (testing for the effect of time period only). For the t-test, the assumption of normality was verified with Shapiro-Wilk tests and the equality of variance with Bartlett’s tests. We checked all GLMM and GLM models for overdispersion using the overdisp_fun function from <https://bbolker.github.io>.

3. Whole genome sequencing and SNP calling

We separated the eggs from the sediment with the sugar flotation method described above. Each pool consisted of 200 eggs of *L. minutus* (except for Lumsden pre-acidification: 155 eggs) that were isolated in 70% ethanol after identification under high-resolution stereomicroscopes (Olympus). The resting eggs were first crushed with a plastic pestle after being flash-frozen with liquid nitrogen. We then extracted DNA from the crushed eggs with the QIAGEN MagAttract HMW kit, following the manufacturer's instructions. DNA quality control (QC) was done with the Agilent gDNA 165 kb kit on the Femto pulse system. Libraries were prepared for Illumina sequencing with the Lucigen NxSeq AmpFREE kit and then had QC performed, then PCR enriched for 6 cycles followed by a final QC. Quality control, library preparation, and sequencing were conducted at the McGill Genome Center.

We trimmed reads with fastp [9] with a minimum base quality score of 20 in a sliding window starting at the 3’ end and a minimum length of 50 bp. The reference-free SNP discovery method implemented in discosnp++ uses a de Bruijn graph analysis of k-mers, produces SNP datasets equivalent to reference-based approaches [10], and has been previously used for a copepod genomic dataset [11]. We used Illumina reads from another *Leptodiaptomus minutus* population from Quebec, Canada (L145, unpublished data) as an anchor for the SNP calling. This library was generated with 200 taxonomically identified pooled *L. minutus* individuals. Briefly, we extracted the DNA using the MagAttract HMW kit (QIAGEN, Toronto, ON, Canada) following manufacturer steps. DNA was run for QC with the Agilent gDNA 165 kb kit on the Femto pulse system. The library was prepared for Illumina sequencing with the Illumina Truseq DNA PCR-free kit and QCed and sequenced with Novaseq 6000, then trimmed with fastp similarly to the libraries generated for the George and Lumsden populations. The trimmed fastq files for the anchor populations are provided in the Dryad Digital Repository (<http://datadryad.org/share/LINK_NOT_FOR_PUBLICATION/e7KFZzHWsT4kJX6yXOnpMLo1AWJKcNY7aYk9mKj1rfw>). We ran discosnp++ with the branching strategy (option -b 1), which allows a good compromise between precision and recall [10], extending polymorphism with left and right unitigs (only one SNP per contig) and a minimum coverage of 5 per pool. We converted the vcf file output by discosnp++ into a "pooldata" object with the poolfstat package in R [12], with a haploid pool size of 400 for all populations (except for the size of 310 for Lumsden pre-acidification), minimum coverage of five per pool, maximum coverage of 300, a minimum read count per base of two, a minimal minor allele frequency of 0.0032 (to remove singletons) and removing indels. We excluded the data for the anchor population L145 for the rest of the downstream analyses after the SNP calling and filtering.

4. Detecting signals of selection

SNP-specific pairwise F_ST_ values were obtained with the compute.pairwiseFST function using the ANOVA method (default) and the same parameters as described in the filtering step above. Because genomic scans using pairwise F_ST_ do not consider population structure and demographic processes, we also used the hierarchical models implemented in Baypass [13], with the core model as an outlier analysis and the auxiliary (Aux) model as an environmental association analysis. This analysis takes the demographic history into account, by correcting for the confounding effects of population structure during the detection of outlier SNPs. This is done by first estimating the scaled covariance matrix Ω of population allele frequencies in the core model, which recapitulates population history. The scaled covariance matrix Ω corresponds to the allele frequency deviation between populations from an allele frequency averaged across populations. Closely related populations tend to have more similar deviations (i.e. covariance) from the average [14]. The scaled covariance matrix Ω is explicitly accounted for during the detection of outlier SNPs [13]. We divided the poolfstat dataset into four pseudo-independent datasets by converting the pooldata object to four Baypass input files, using the "thinning" subsampling method with a sub-sample size of 842,255 SNPs. We ran both the core and auxiliary models with default parameters. We first used the core model to estimate the scaled covariance matrix of population allele frequencies Ω (Fig. S2). This model also allows the estimation of the XtX statistics, which is equivalent to SNP-specific F_ST_ and is based on the variance in allele frequencies between all populations [15], accounting for Ω and derived p-values under a χ^2^ distribution with 5 degrees of freedom (N = 5 pools, bilateral test). We confirmed that the p-values behaved well based on the shape of their histogram (Fig. S6, [13]). We selected the Aux model for the environmental association analysis as it is suitable for analyses of time samples such as experimental evolution datasets (see Baypass manual). This model computes the regression coefficient β_ik_ of the association between the SNP allele frequencies and the covariable (here positive and negative association with the acidification), as well as an associated Bayesian auxiliary variable δ_ik_ from which a Bayes factor BF_mc_ is derived. The acidification covariable was standardized to $\hat{\mu}$ = 0 and $\hat{\alpha^{2}}$ = 1 as recommended. We compared the four Ω matrices visually to ensure that the results were concordant between sub-datasets, then we combined the statistics estimated for each SNP.

5. Population structure and demography.

We used the diffusion approximation approach from δaδi to identify changes in effective population size N_e_ [16]. This approach was selected to reconstruct demographic changes because copepod exoskeletons do not preserve well over time in sediment archives and thus cannot be used as a measure of past abundance [17]. The most complex demographic model (“bottlegrowth”) analyzed with δaδi has an ancestral effective population size *N_ref_*, then a genetic bottleneck at time T_B_ (scaled time between the bottleneck and present) corresponding to a decrease in effective population size toward an effective population of size *NuB*, followed by exponential recovery in N_e_ to reach the effective population size *NuF*. As we knew the approximate time of the potential bottleneck (Lumsden = 1976, set as 42 years before sampling; George = 1972, set as 46 years before sampling; with two generations per year [18]), T_B_ was defined as a fixed parameter, and the parameter $\theta=4\mu L$ as an explicit parameter, for all models except the neutral model. We defined *θ* with *μ* the mutation rate of $2.64{\times10}^{-9}$ substitutions per site per year from the snapping shrimp *Alpheus spp.* [19] and *L* the effective sequenced length of 319,822,866 bp for Lumsden and 334,764,074 bp for George, calculated as:

L ≈ total length of sequence analyzed $\times$ SNPs retained for use in δaδi / total SNPs in analyzed sequence.

To generate an allele frequency table, we used the summary_yij_ypij.out file output from Baypass with the Core model and selected the M_P column corresponding to the mean of the posterior distribution of the α_ij_ parameter, which is closely related to the allele frequency of the reference allele (chosen arbitrarily). We filtered out SNPs that were detected as outliers (putatively under selection) in the Baypass (core and Aux models) and F_ST_ scan analyses, and SNPs with an allele frequency < 0.0025, which corresponds to an allele count of 0, thus uninformative in δaδi. The filtering retained 3,681,690 SNPs for Lumsden and 3,853,688 SNPs for George. To transform the allele frequency data into the input format of δaδi, we used the dadi_input_pools function from the genomalicious R package [20] applying the “probs” parameter in the methodSFS option. As we lacked information on the ancestral allele state, we inferred the folded SFS with δaδi. We used the default local optimizer on the log of parameters and optimized the parameters for each model until convergence was reached (three runs falling within 1% of the best likelihood). We compared the two nested models (bottlegrowth and growth) with an adjusted likelihood ratio test, and the two remaining models (two epochs and neutral) based on the large differences in the likelihoods and residuals between these models and the model showing the highest likelihood (bottlegrowth and growth). As we obtained unlikely results during the conversion of the parameters of the best model for each population, possibly due to imprecise mutation rates from a distantly related species, we do not report the results of parameter conversion. To obtain the uncertainties on the parameters while accounting for the effect of linkage, we used bootstrapping and the Godambe Information Matrix approach [21]. For this, we generated 100 bootstrapped datasets with a chunk size of $1\times{10}^{5}$ bp. Note that due to the SNPs being called from contigs in the reference-free SNP discovery approach, the bootstrapping was executed with individual SNPs, which possibly led to the underestimation of uncertainties.

**Table S1: Environmental characteristics of the study sites (June 2018-2019)**

| Lake | Maximum depth (m) | Surface temperature | pH | Conductivity (µS/cm) | Dissolved organic carbon (mg/L) | Total Phosphorus (µg/L) | Total Nitrogen (mg/L) |
| --- | --- | --- | --- | --- | --- | --- | --- |
| George | 36.5 | 25.4 | 7.05 | 18.9 | 2.7 | 3.2 | 0.20 |
| Lumsden | 25 | 25.0 | 6.13 | 11.1 | 2.5 | 3.4 | 0.18 |

| Time period | Lake | Depth (cm) | Number of cores pooled | pH treatment | Number of replicates | Average number of individuals $\pm$SD |
| --- | --- | --- | --- | --- | --- | --- |
| Pre-acidification | George | 9-12 | 32 | Acidic | 0 | NA |
|  |  |  |  | Neutral | 0 | NA |
|  | Lumsden | 10-13 | 32 | Acidic | 5 | 20 |
|  |  |  |  | Neutral | 5 | 19.8$\pm$0.4 |
| Acidification | George | 5-8 | 32 | Acidic | 5 | 19$\pm$2.2 |
|  |  |  |  | Neutral | 2 | 19.5$\pm$0.7 |
|  | Lumsden | 4-7 | 30 | Acidic | 5 | 20 |
|  |  |  |  | Neutral | 5 | 20.2$\pm$0.4 |
| Recovery | George | 0-2 | 27 | Acidic | 5 | 20 |
|  |  |  |  | Neutral | 3 | 18.3$\pm$2.9 |
|  | Lumsden | 0-2 | 26 | Acidic | 5 | 20 |
|  |  |  |  | Neutral | 5 | 19.2$\pm$1.1 |

**Table S2: Details of the resurrection ecology experimental design**

**Table S3: Results of the analysis of survival, fecundity, and development times with generalized linear mixed effect models (GLMM), generalized linear models (GLM), and t-tests.** Lifetime survival indicates survival from hatching to adulthood and early survival from hatching to copepodid stage. For the GLMMs, a ΔAIC > 2 indicates that the effect is significant. Significant fixed effects are indicated in bold.

| **Response variable** | **Lake** | **Type model** | **Test** | **Fixed effects** | **df** | **p-value**  **ΔAIC** |
| --- | --- | --- | --- | --- | --- | --- |
| **Lifetime survival** | Lumsden | GLMM binomial logit | AICc | Time period $\boldsymbol{\times}$ pH | 5 | -2.95 |
|  |  |  |  | Time period | 3 | 10.83 |
|  |  |  |  | pH | 4 | 2.01 |
| **Early survival** | Lumsden | GLMM binomial logit | AICc | Time period $\boldsymbol{\times}$ pH | 5 | -5.9 |
|  |  |  |  | Time period | 3 | 8.41 |
|  |  |  |  | pH | 4 | -0.90 |
| **Lifetime survival** | George | GLMM binomial logit | AICc | Time period $\boldsymbol{\times}$ pH | 4 | -1.87 |
|  |  |  |  | Time period | 3 | 7.55 |
|  |  |  |  | pH | 3 | 3.48 |
| **Early survival** | George | GLMM binomial logit | AICc | Time period $\boldsymbol{\times}$ pH | 4 | - 4.44 |
|  |  |  |  | Time period | 3 | 5.94 |
|  |  |  |  | pH | 3 | -3.23 |
| **Fecundity** | Lumsden | GLM quasi-poisson | Quasi-F | Time period $\boldsymbol{\times}$ pH | 24 | 0.20 |
|  |  |  |  | Time period | 26 | 0.04 |
|  |  |  |  | pH | 28 | 0.38 |
| **Fecundity** | George | GLM quasi-poisson | Quasi-F | Time period $\boldsymbol{\times}$ pH | 11 | 0.08 |
|  |  |  |  | Time period | 12 | 0.07 |
|  |  |  |  | pH | 13 | 0.15 |
| **Lifetime development** | Lumsden | GLM gamma log | LRT | Time period $\boldsymbol{\times}$ pH | 17 | 0.78 |
|  |  |  |  | Time period | 20 | 5.59$\boldsymbol{\times}\boldsymbol{10}^{\boldsymbol{-7}}$ |
|  |  |  |  | pH | 19 | 0.93 |
| **Early development** | Lumsden | GLM gamma log | LRT | Time period $\boldsymbol{\times}$ pH | 22 | 0.16 |
|  |  |  |  | Time period | 25 | 0.002 |
|  |  |  |  | pH | 24 | 0.39 |
| **Lifetime development** | George | - | t-test | Time period | 4 | 0.121 |
| **Early development** | George | - | t-test | Time period | 12 | 0.07 |

**Table S4: Fixed effect coefficients of the GLMs and GLMMs and their confidence intervals after conversion with an exponential function for each response variable and lake.** The converted coefficients are given as the first level compared to the second (reference) level. The relationship is significant if the converted coefficients do not include 1 (in bold), is positive if > 1, and negative if < 1. When the converted coefficient is < 1, the inverse is given in parentheses and indicates how much less likely an event is to be observed. For the t-tests results (development time for George), the estimate represents the difference (in days) between time periods.

| **Response variable** | **Lake** | **Fixed effect coefficients** | **Comparison** | **Estimate** | **Lower limit** | **Upper limit** |
| --- | --- | --- | --- | --- | --- | --- |
| **Lifetime survival** | Lumsden | Odds ratio | Acidification vs pre-acidification | 7.98 | 2.96 | 21.51 |
|  |  |  | Recovery vs pre-acidification | 1.31 | 0.46 | 3.72 |
|  |  |  | Neutral pH vs acidic pH | 2.59 | 1.14 | 5.87 |
| **Early survival** | Lumsden | Odds ratio | Acidification vs pre-acidification | 3.36 | 1.42 | 7.97 |
|  |  |  | Recovery vs pre-acidification | 0.59 (1.69) | 0.25 | 1.41 |
|  |  |  | Neutral pH vs acidic pH | 1.70 | 0.84 | 3.45 |
| **Lifetime survival** | George | Odds ratio | Recovery vs acidification | 0.12 (8.33) | 0.04 | 0.40 |
|  |  |  | Neutral pH vs acidic pH | 5.35 | 1.66 | 17.19 |
| **Early survival** | George | Odds ratio | Recovery vs acidification | 0.27 (3.70) | 0.11 | 0.67 |
|  |  |  | Neutral pH vs acidic pH | 1.45 | 0.57 | 3.75 |
| **Fecundity** | Lumsden | Incident rate ratios | Acidification vs pre-acidification | 23.0 | 1.32 | 9.30$\boldsymbol{\times}\boldsymbol{10}^{\boldsymbol{5}}$ |
|  |  |  | Recovery vs pre-acidification | 9.5 | 0.36 | 3.96$\boldsymbol{\times}\boldsymbol{10}^{\boldsymbol{5}}$ |
| **Fecundity** | George | Incident rate ratios | Recovery vs acidification | 0.32 (3.13) | 0.05 | 1.41 |
| **Lifetime development** | Lumsden | Fold change | Acidification vs pre-acidification | 0.67 (1.49) | 0.58 | 0.77 |
|  |  |  | Recovery vs pre-acidification | 0.73 (1.36) | 0.61 | 0.86 |
| **Early development** | Lumsden | Fold change | Acidification vs pre-acidification | 0.82 (1.22) | 0.72 | 0.93 |
|  |  |  | Recovery vs pre-acidification | 0.98 (1.02) | 0.87 | 1.13 |
| **Lifetime development** | George | - | Recovery vs acidification | 4.34 | - | - |
| **Early development** | George | - | Recovery vs acidification | 1.68 | - | - |

**Table S5: Demographic analysis with dadi of Lumsden and George recovery samples for four models (genetic bottleneck followed by growth “bottlegrowth”, genetic bottleneck only “two epochs”, growth and neutral).** Models are presented according to their log likelihood (log L), with the number of parameters in each model (k). For the best models (bottlegrowth and growth, underlined), we show the estimated scaled parameters *NuB* (ratio of effective population size after the bottleneck to ancestral N_e_), *NuF* (ratio of N_e_ after the recovery to ancestral N_e_), *Nu* (ratio of contemporary N_e_ to to ancestral N_e_) and *Theta* ($\theta=4\mu L$, with *μ* the mutation rate and *L* the effective sequenced length; see main text), with the upper and lower bounds of the 95% confidence interval shown in brackets.

| **Population** | **Model** | **Log likelihood** | ***k*** |  | ***NuB*** | ***NuF*** | ***Nu*** | ***Theta*** |
| --- | --- | --- | --- | --- | --- | --- | --- | --- |
| **George** | Bottlegrowth | -51,561 | 3 |  | $3.5\times{10}^{-5}$ | 6.2 | - | $1.5\times{10}^{6}$ |
|  | Two epochs | -59,672 | 2 |  | - | - | $4.6\times{10}^{-4}$ | $1.5\times{10}^{6}$ |
|  | Growth | -17,690 | 2 |  | - | - | $3.6\times{10}^{-5}$  [-5$.7\times{10}^{-2}$- 5$.7\times{10}^{-2}$] | $2.9\times{10}^{6}$  [2$.9\times{10}^{6}$- $2.9\times{10}^{6}$] |
|  | Neutral | -48,271,793 | 1 |  | - | - | - | 580,578 |
| **Lumsden** | Bottlegrowth | -79,544 | 3 |  | $2.3\times{10}^{-5}$  [-0.3 – 0.3] | 31.9  [31.0 – 32.9] | - | $1.5\times{10}^{6}$  [1$.5\times{10}^{6}$- $1.5\times{10}^{6}$] |
|  | Two epochs | -98,971 | 2 |  | - | - | $3.7\times{10}^{-4}$ | $1.4\times{10}^{6}$ |
|  | Growth | -43,615 | 2 |  | - | - | $2.7\times{10}^{-5}$  [-9$.7\times{10}^{-2}$- 9$.7\times{10}^{-2}$] | $3.0\times{10}^{6}$  [1$.0\times{10}^{6}$- $2.3\times{10}^{6}$] |
|  | Neutral | -45,699,817 | 1 |  | - | - | - | 551,856 |

**Table S6: Pairwise F_ST_ matrix between the 5 temporal samples.** LUM: Lumsden Lake, GEO: George Lake, PRE: pre-acidification, ACID: acidification, REC: recovery.

|  | LUM-PRE | LUM-ACID | LUM-REC | GEO-ACID | GEO-REC |
| --- | --- | --- | --- | --- | --- |
| LUM-PRE |  | 0.03 | 0.02 | 0.08 | 0.09 |
| LUM-ACID |  |  | 0.03 | 0.10 | 0.10 |
| LUM-REC |  |  |  | 0.09 | 0.10 |
| GEO-ACID |  |  |  |  | 0.03 |
| GEO-REC |  |  |  |  |  |

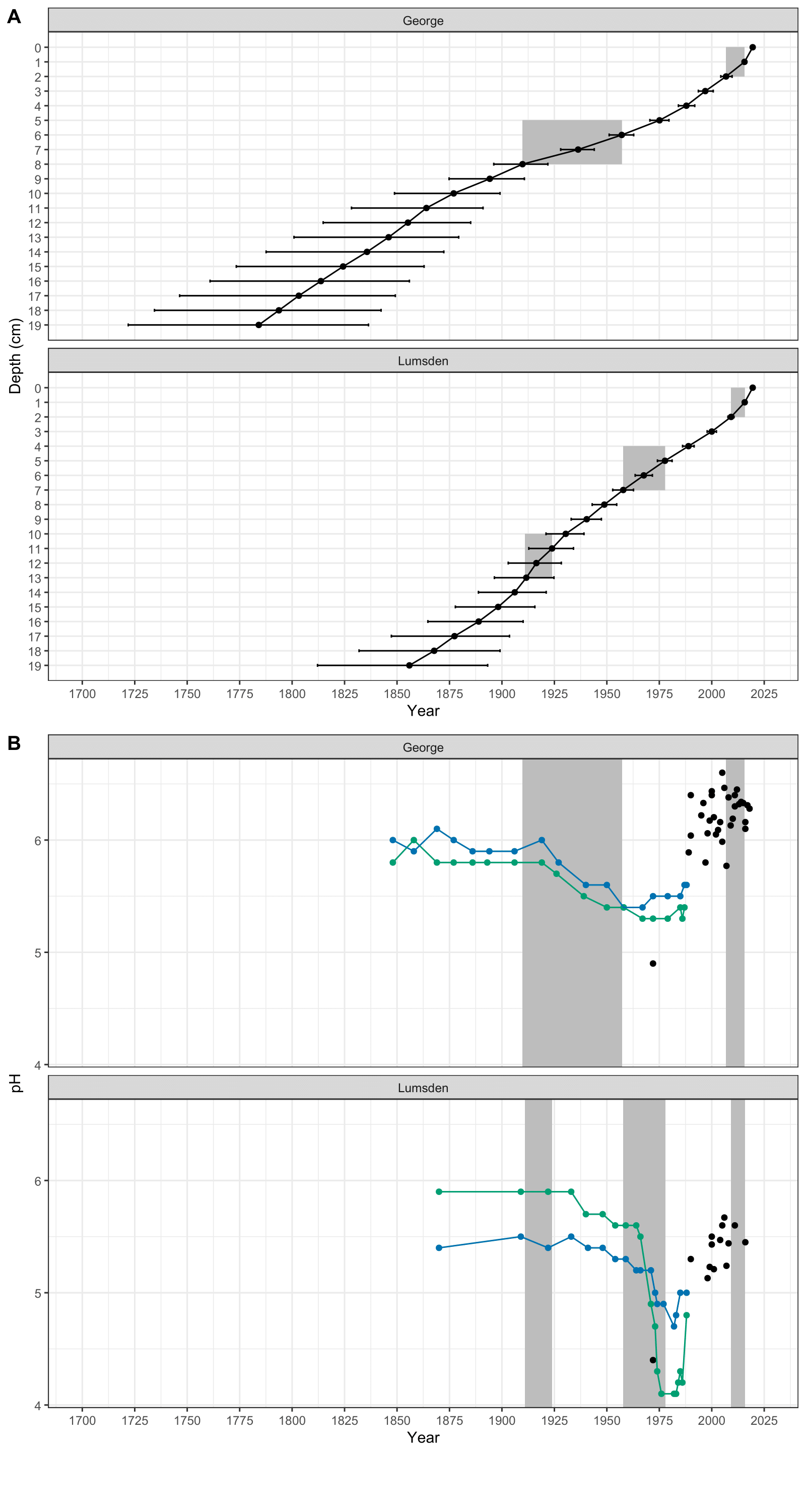
**Figure S1. ^210^Pb-dating of sediment cores and historical pH trajectory for George and Lumsden lakes**. **A:** ^210^Pb sediment core dating reconstructed using the rplum package. Grey boxes indicate the selected depth sections and start/end date for each time period (chronologically: pre-industrial, acidification, recovery). Dots are the mean date for each section, and errors bars shows the minimum and maximum dates. **B:** Historical lake pH based on reconstruction from diatom and chrysophyte paleofossils [22], and contemporary lake pH as measured by the Ontario Ministry of the Environment and from the literature [7] for George and Lumsden lakes. Grey boxes approximate the start and end date of the time periods (pre-industrial, acidification, recovery).

**Fig S2: Population genetic structure and demographic history.** Heatmap of the scaled covariance matrix Ω (with ρ_ij_ the correlation coefficient between pairs of populations) with hierarchical clustering tree (using the average agglomeration method), obtained from the core model of Baypass.

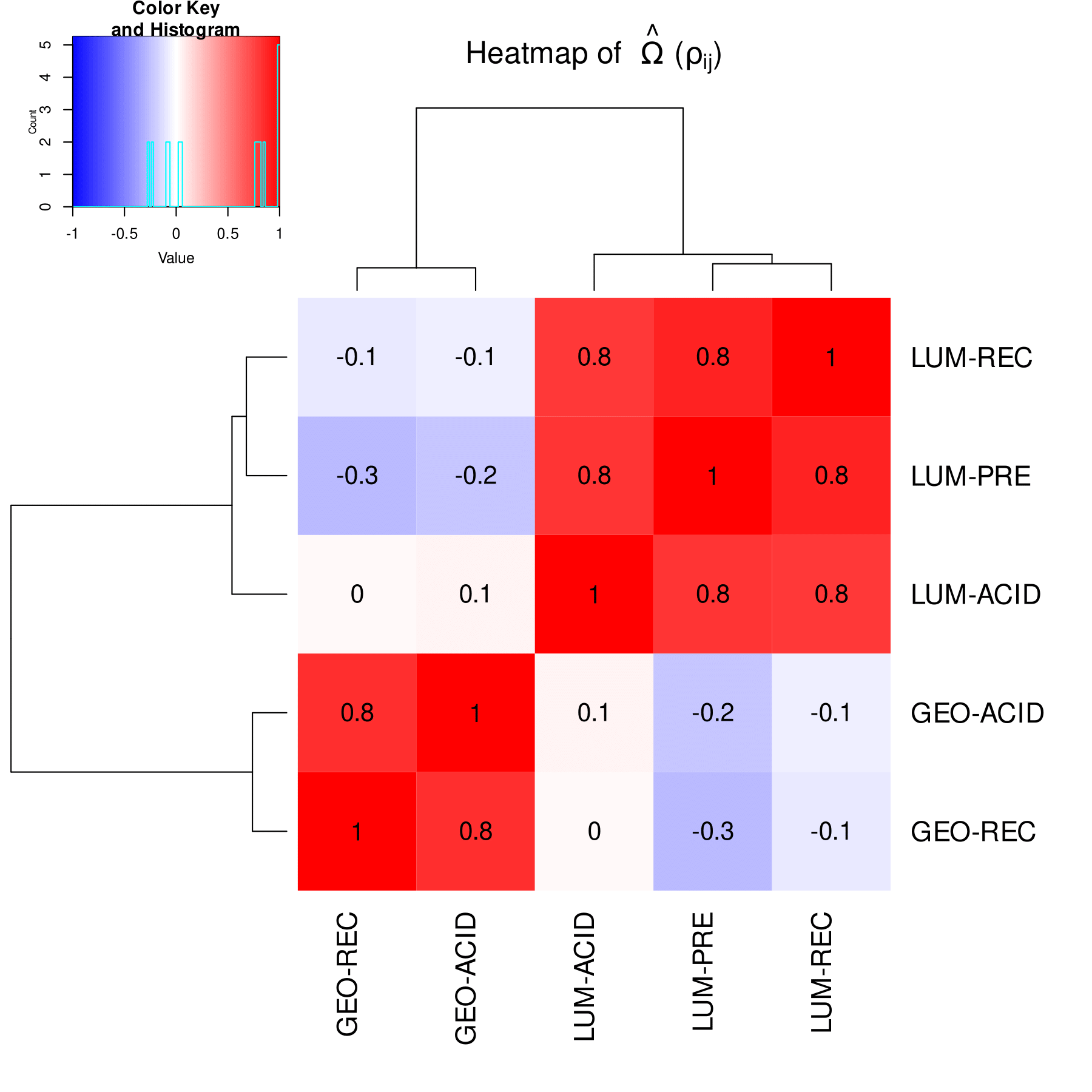

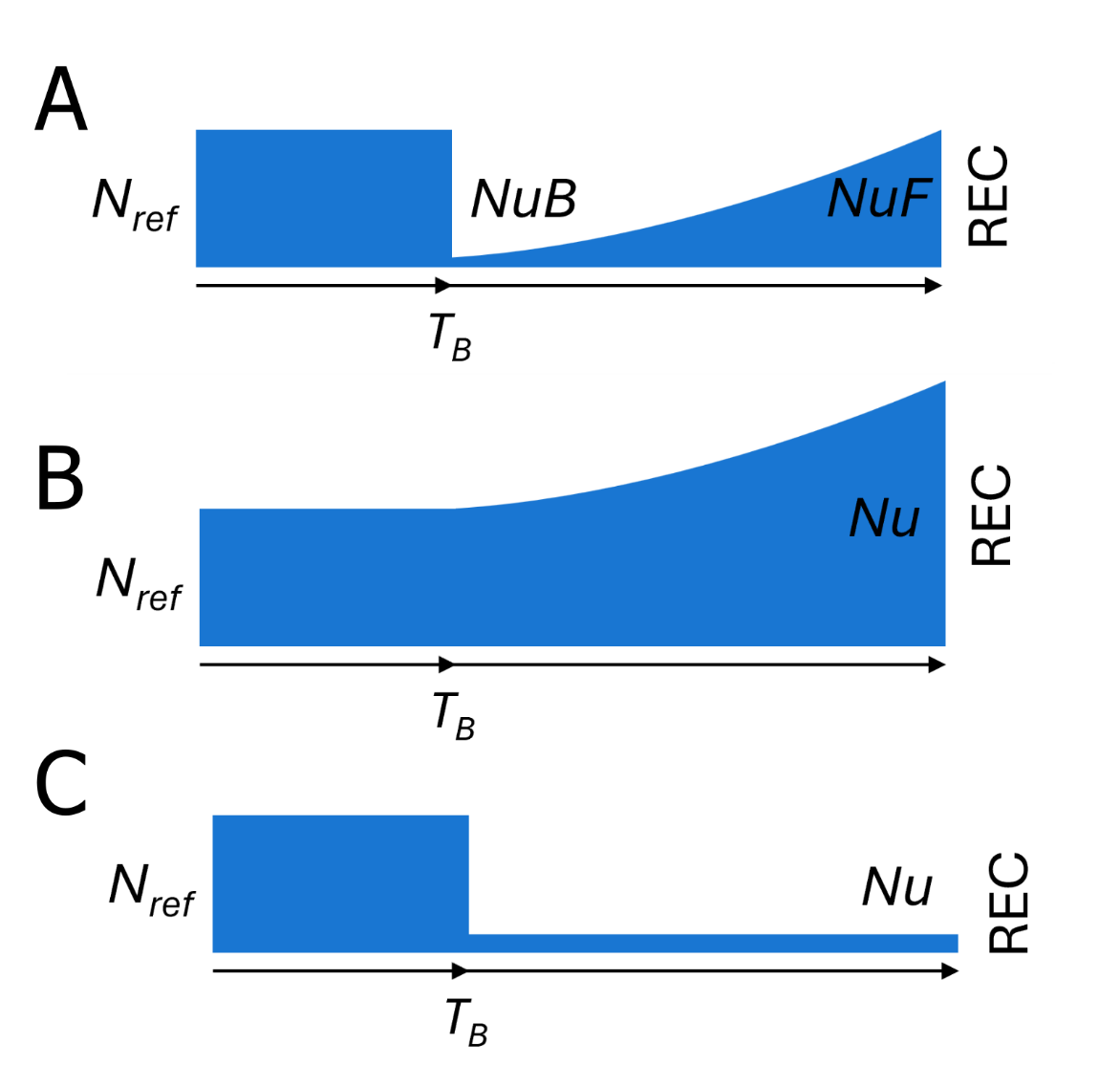
**Figure S3: Alternative demographic model tested with dadi.** Visual 1D demographic model of a bottleneck without N_e_ recovery (two-epochs) after the time of acidification T_B_ (scaled time between the bottleneck and the present). The parameter *Nu* is the ratio of the effective population size after the bottleneck to the ancestral effective population size *N_ref_*..

**Figure S4: Results of the resurrection ecology experiment for George and Lumsden lakes as a function of time periods of origin for lake acidification history and pH treatment for early development stages (from hatching as nauplii to metamorphosis into copepodid). (A)** Early survival, **(B)** early development time. Each dot represents a measurement for one replicate; the black horizontal bars are the mean responses for each treatment.

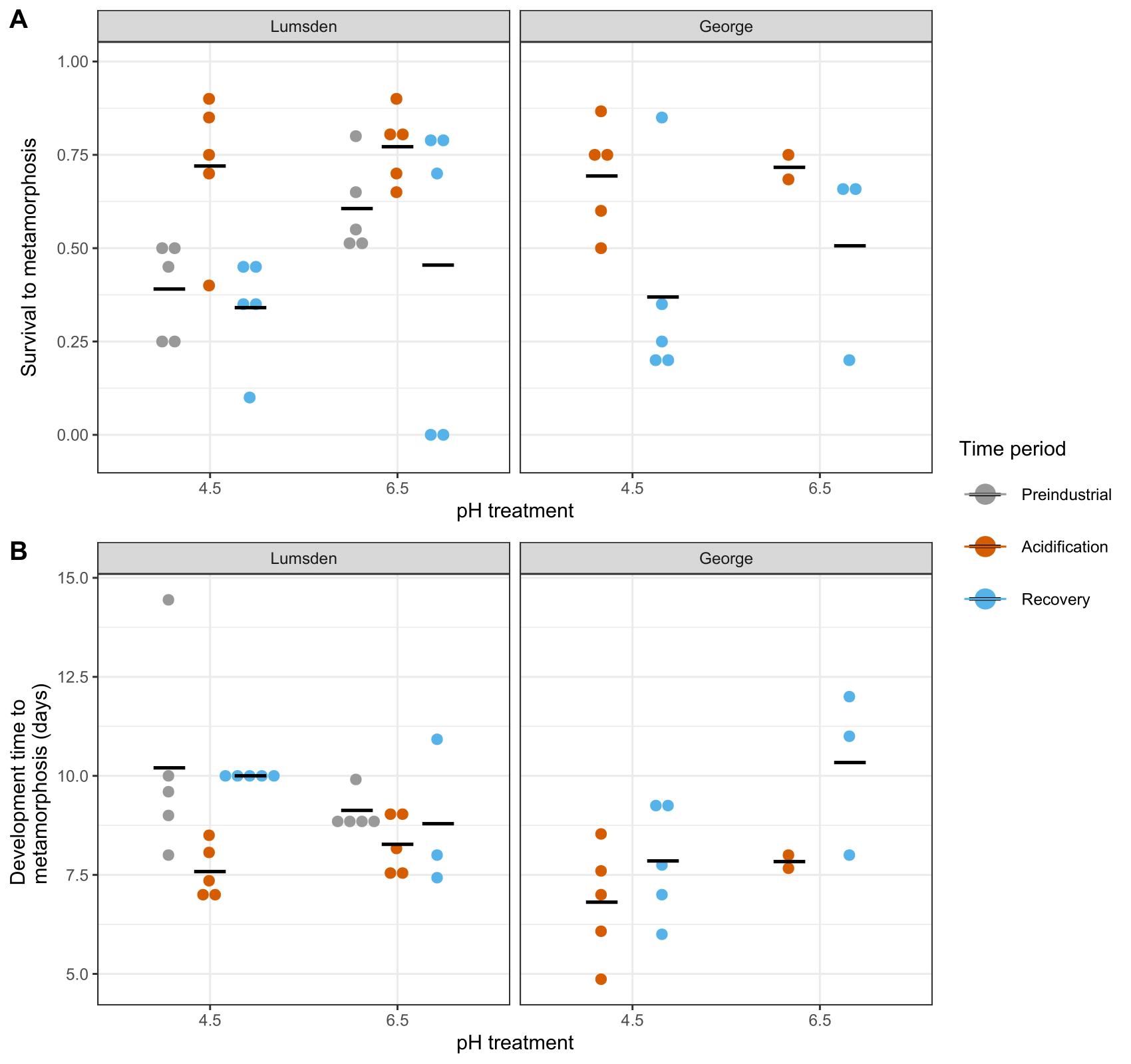

**Figure S5: Alternative growth model tested with dadi for Lumsden Lake copepods from the recovery period**. Top: Folded site frequency spectrum (SFS) for a sample size of 400 (two times the number of individuals) for the observed data (blue line) and the model (red line), showing the logarithm of the number of sites as a function of a given read count. Bottom: residuals of the normalized differences between the observed data and the model.

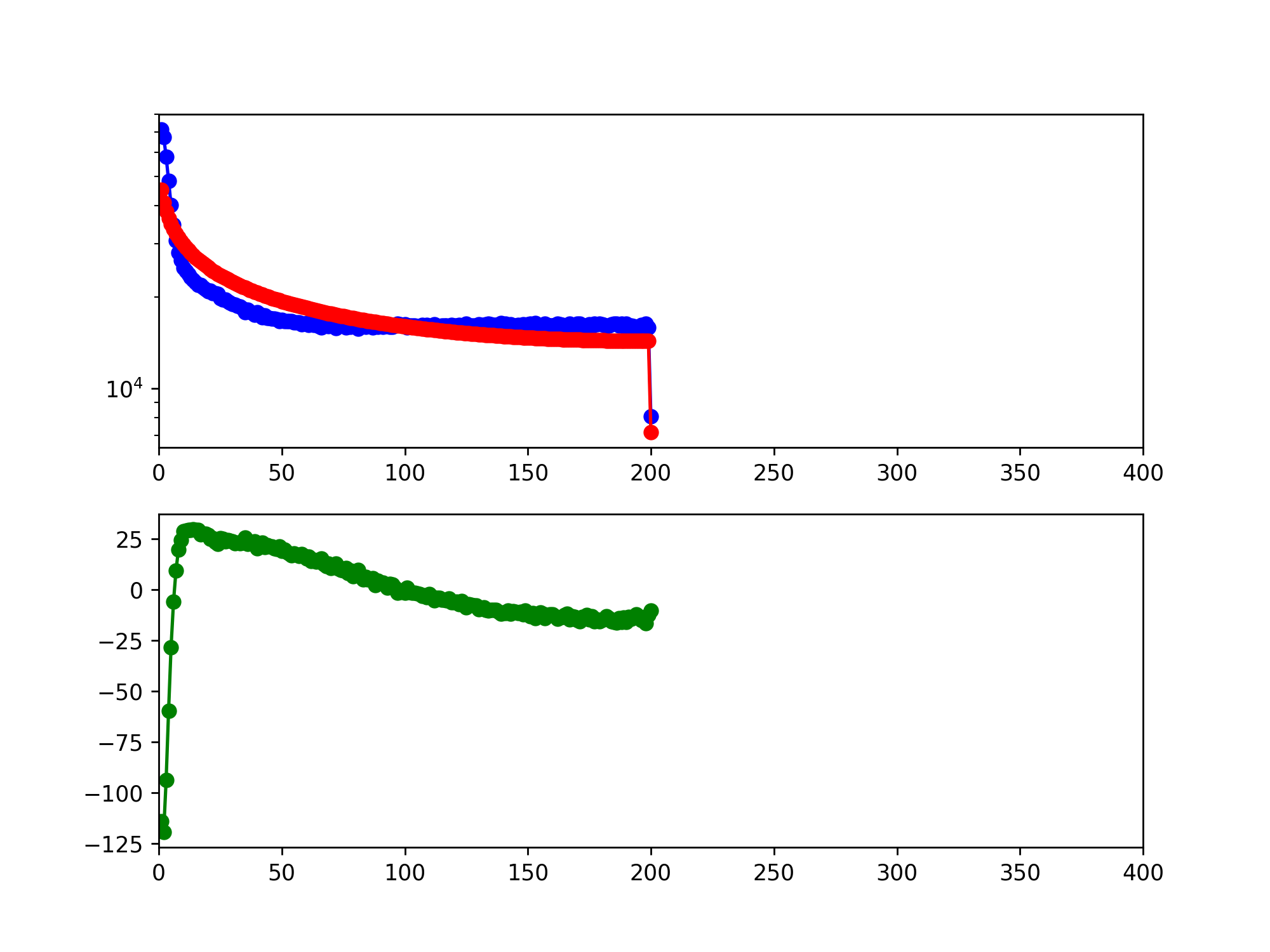

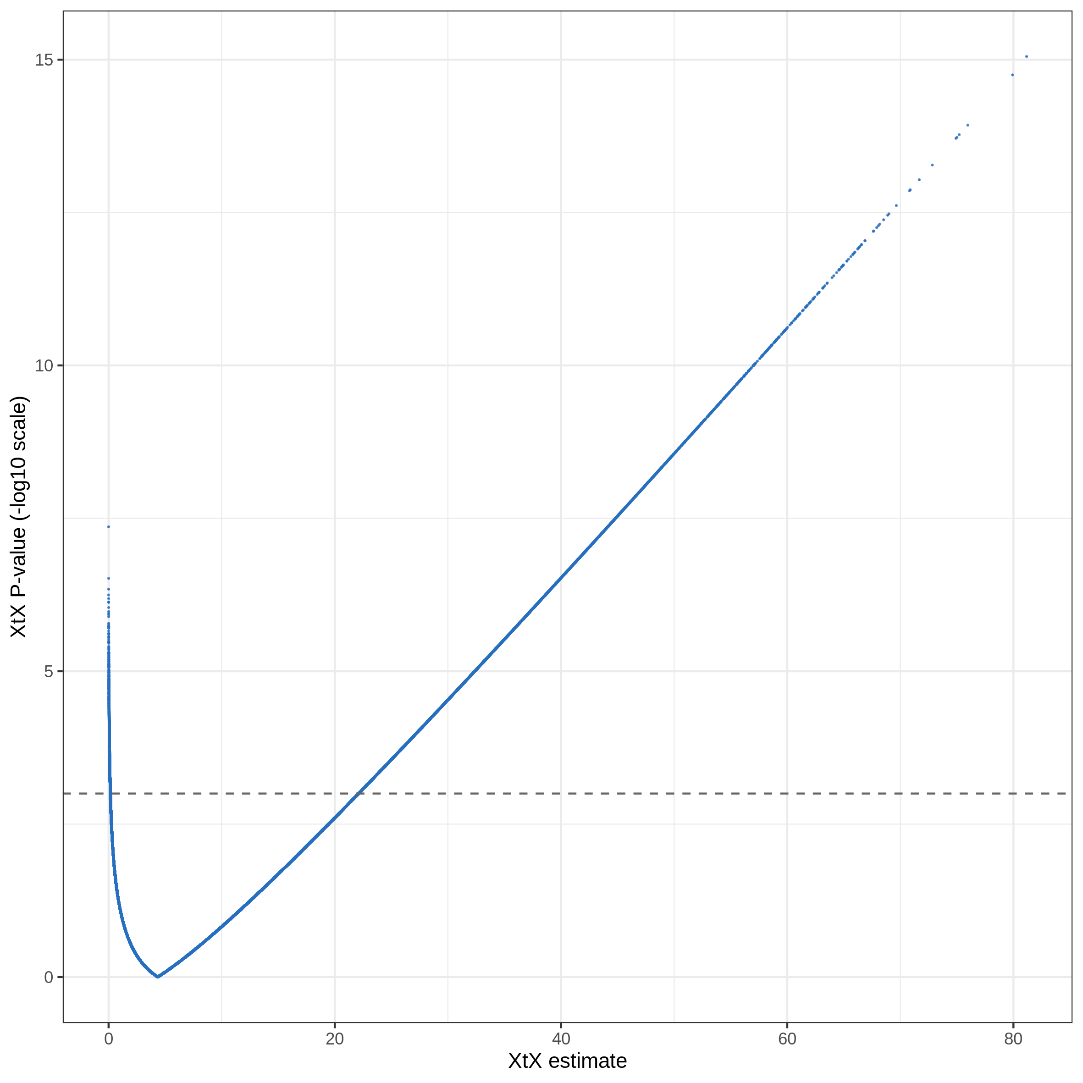

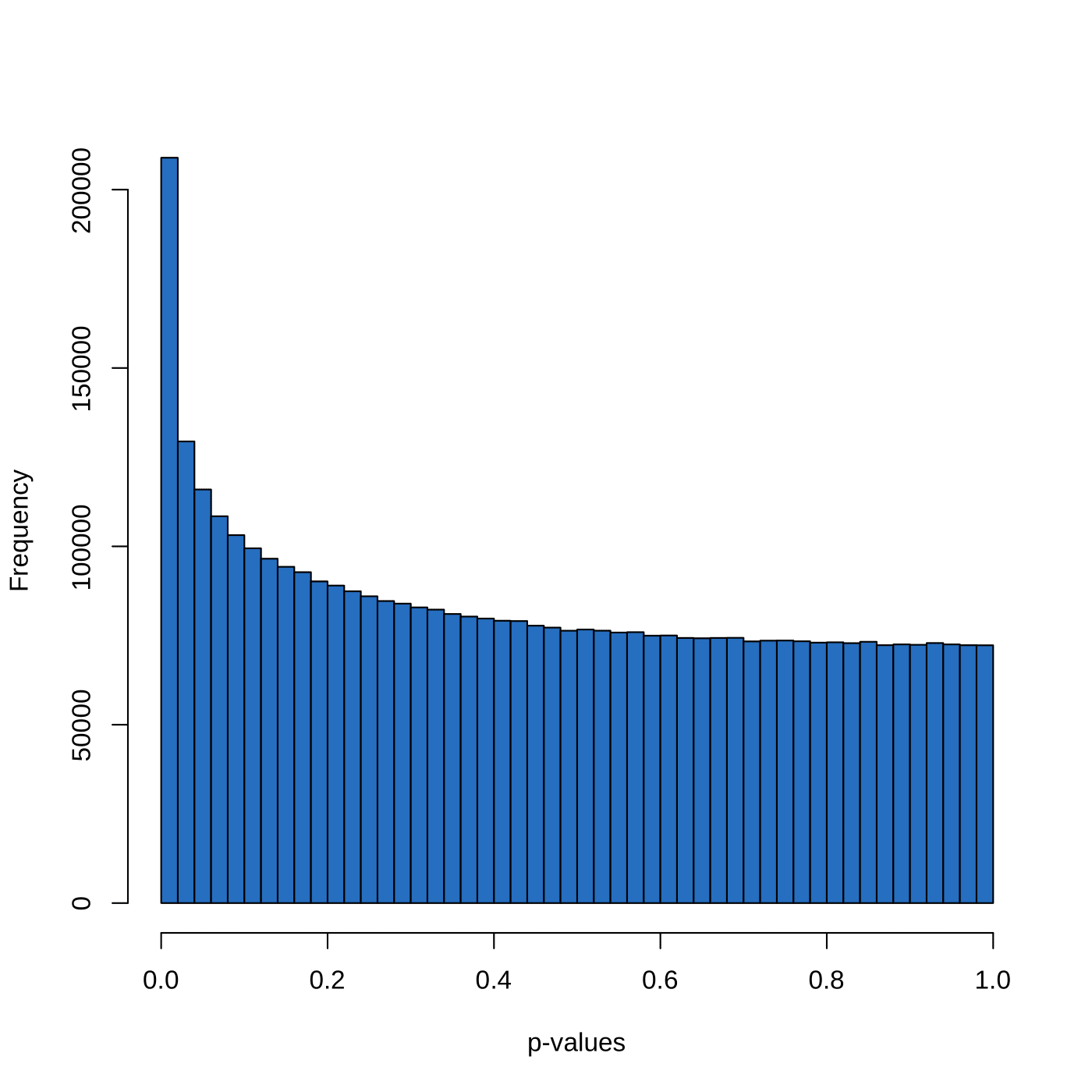
**Figure S6: Outliers SNPs identified with the core model in Baypass.** Left: Histogram of the p-values derived from the XtX estimate. Right: P-values associated with the XtX statistic (on -log10 scale) as a function of the XtX estimate. The horizontal line indicates the threshold of p-values < 0.001.

**Figure S7: Venn diagram of the outliers detected with the core and auxiliary models in Baypass in common with the outliers of the FST scan for George (acidification vs recovery comparison) and Lumsden (outliers in at least one of the temporal comparisons: pre-acidification vs acidification and acidification vs recovery).** Outlier SNPs shared by the different analyses are shown in the shaded areas.

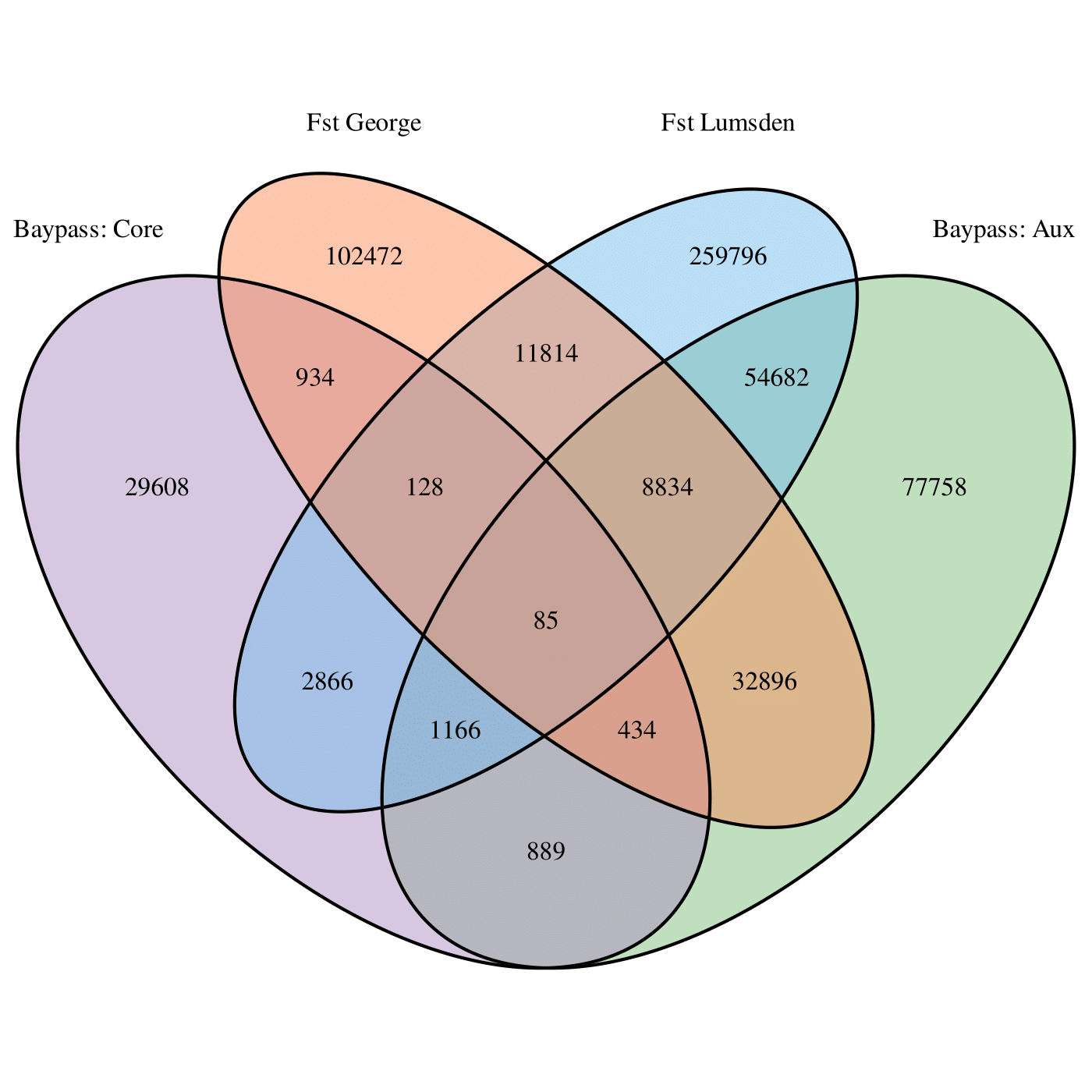

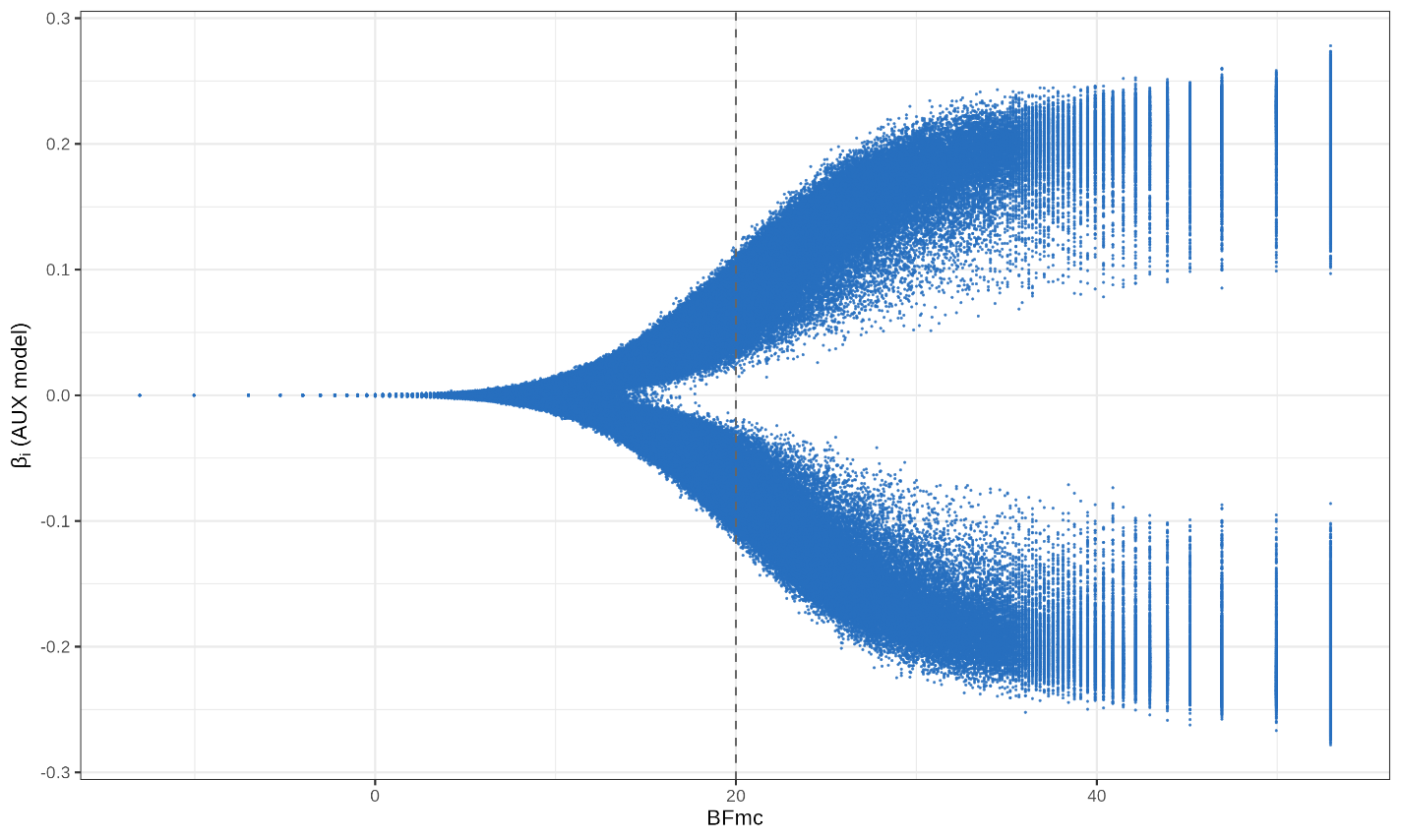
**Fig S8: Correlation coefficient β_i_ between allele frequencies and the acidification as a function of Bayes Factor in Baypass**. SNPs were considered as outliers when BFmc > 20
